# Lineage-specific intolerance to oncogenic drivers restricts histological transformation

**DOI:** 10.1101/2023.06.21.545980

**Authors:** Eric E. Gardner, Ethan M. Earlie, Kate Li, Jerin Thomas, Melissa J. Hubisz, Benjamin D. Stein, Chen Zhang, Lewis C. Cantley, Ashley M. Laughney, Harold Varmus

## Abstract

Lung adenocarcinoma (LUAD) and small cell lung cancer (SCLC) are thought to originate from different epithelial cell types in the lung. Intriguingly, LUAD can histologically transform into SCLC following treatment with targeted therapies. Here we designed models to follow the conversion of LUAD to SCLC and found the barrier to histological transformation converges on tolerance to Myc, which we implicate as a lineage-specific driver of the pulmonary neuroendocrine cell. Histological transformations are frequently accompanied by activation of the Akt pathway. Manipulating this pathway permitted tolerance to Myc as an oncogenic driver, producing rare, stem-like cells, transcriptionally resembling the pulmonary basal lineage. These findings suggest histological transformation may require the plasticity inherent to the basal stem cell, enabling tolerance to previously incompatible oncogenic driver programs.

**One-Sentence Summary:** By modeling histological transformation of lung cancer, we uncover neuroendocrine-specific tolerance to Myc as an oncogenic driver.

## Main Text

Histological transformation (abbreviated hereafter as “HT”) is a poorly understood process whereby a cancer’s initial histology is altered and presents as a new histologic type of cancer. These changes are presumed to be under selective pressure of an oncogene-targeted therapy. Best described in the context of EGFR inhibition in lung adenocarcinoma (LUAD) (*1–3*) and androgen receptor inhibition of prostate adenocarcinoma (*4, 5*), these transformations most commonly lead to neuroendocrine and/or squamous differentiation (*6*). They are considered “off-target” forms of acquired resistance as the cancer’s proliferative character is no longer dependent on the original oncogenic driver pathway. In the earliest reports of HT, it was unclear whether the recurrent tumors were independent primary SCLCs, sub-clonal selection under the pressure of targeted therapy or products of the direct conversion of LUAD into SCLC. Supporting evidence for direct conversion relies heavily on the continued presence of a LUAD driver oncogene within a cancer that is histologically SCLC. However, oncogenic mutations in the genes that commonly drive the formation of LUADs have rarely been encountered in genome sequencing studies of primary, treatment-naïve SCLC (*7, 8*).

Critically, some post-transformation samples have shared, clonally related mutations present in the LUAD prior to treatment with targeted therapy; however, protein expression of the LUAD driver is conspicuously absent post-transformation (*9*). A consistent requirement for HT is the loss of the RB tumor suppressor (*4, 10*) and EGFR-driven LUADs with pre-existing inactivation of *TP53* and *RB1* are at an especially high risk for undergoing HT (*1, 11*).

Primary LUAD and SCLC are thought to develop from distinct cell types in the lung: the alveolar type II (AT2) cell and the pulmonary neuroendocrine cell (PNEC), respectively. Much is known about surfactant-producing AT2 cells as a cell of origin for LUAD, which the epidermal growth factor (EGF) signaling pathway is an established mitogenic program (*12–14*). Hence, activating mutations and amplifications of genes that encode proteins in the mitogen activated protein kinase (MAPK) pathway, including *RAS* and *EGFR* are common in LUAD (*15*). In contrast, SCLC is thought to arise predominantly from pulmonary neuroendocrine cells (PNECs) (*8, 16, 17*). PNECs are rare, found near anatomic branchpoints of the large airway and function as sentinels for inhaled pathogens and environmental changes – signaling to other cells and tissues through both electrochemical innervation and secretory function (*17–19*). Due to their scarcity, much less is known about PNEC biology, including what signaling events drive their proliferation.

We hypothesized that the complexity of HT may be simplified to represent a mechanism by which a cell can change its oncogenic driver program. We set out to address several fundamental unknowns: (i) how an adenocarcinoma transforms to a high-grade neuroendocrine cancer, (ii) what the intermediate steps in HT are, and (iii) what the oncogenic driver program of the neuroendocrine lineage is. Here we address these questions by combining models of genetically engineered lung tumorigenesis, lineage tracing and chronologic single cell RNA-sequencing to map cell type-specific oncogenic driver programs and their contribution to HT. The data derived from these approaches converge on the concept of lineage-dependent tolerance of driver oncogenes, especially a unique cellular tolerance for Myc as the oncogenic driver in the PNEC lineage.

## Results

### Distinct tumor histologies in an isogenic mouse model

To chronicle HT we generated a new genetically engineered mouse model that combines the conditional expression of *Myc*, *rtTA3* and *tdTomato* (*lox-stop-lox* alleles) with the loss of tumor suppressors *Rb1* and *Trp53 (floxed* alleles) and a doxycycline (DOX)-inducible, oncogenic EGFR transgene (*20*). The model (herein abbreviated “ERPMT”) allows for two classes of manipulations: (i) control of the cell of origin, through lineage-restricted expression of Cre recombinase using adenoviral vectors with cell type-specific promoters (*21*), and (ii) oncogenic EGFR^L858R^ expression, controlled in a DOX-dependent manner (**Fig. 1A**). Tumors initiated in the AT2 or PNEC lineages in ERPMT mice produced histologically distinct lung tumors with opposing dependencies on the presence or absence of DOX. If expression of Cre was initiated in the AT2 lineage and the mice received DOX, the ERPMT model developed an aggressive LUAD. In contrast, no mice off DOX succumbed to disease in this timeframe. In mice receiving DOX, we observed multifocal, glandular lesions throughout all lobes of the airway consistent with LUAD, showing homogenous expression of tdTomato (tdTom) and lack of the neuroendocrine marker synaptophysin (**Fig. 1B**). Conversely, if expression of Cre was initiated in the PNEC lineage in the ERPMT model, then the cohort off DOX developed an aggressive, neuroendocrine SCLC and mice on DOX appeared healthy at a time when 100% of the cohort off DOX was moribund (**Fig. 1C**).

**Figure 1.**
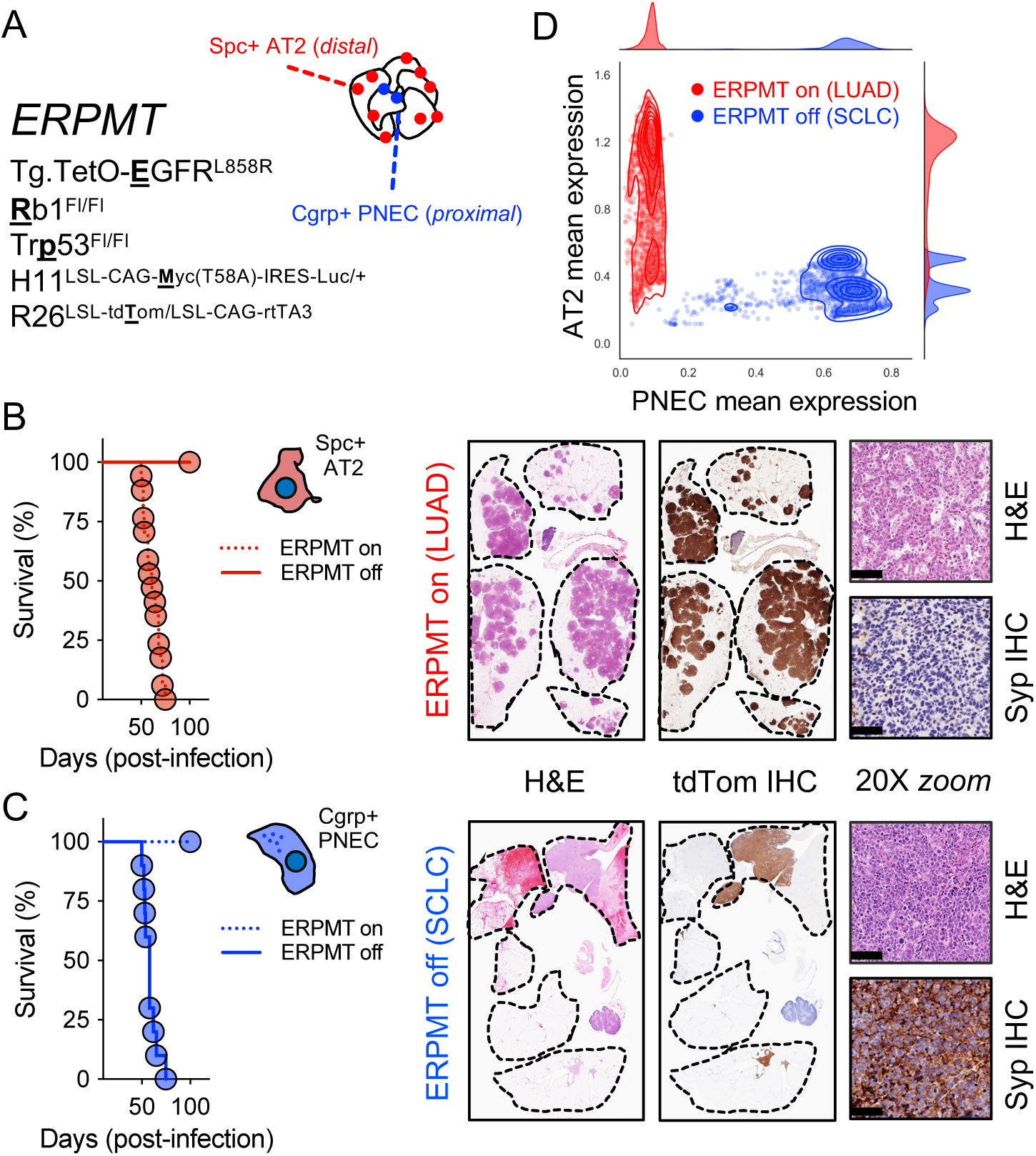
A GEMM model to generate distinct histologic subtypes of lung cancer. A) Nomenclature used for the ERPMT model with abbreviated alleles bolded/underlined and general strategy to use promoter-restricted adenovirus to initiate tumorigenesis in specific airway cells; B) Survival and histologic appearance of the ERPMT model initiated in alveolar type 2 cells (AT2; *red cartoon cell*) or C) pulmonary neuroendocrine cells (PNEC; *blue cartoon cell*) when mice are on doxycycline (DOX) containing (dashed line; n = 10 per group) or control diet (*solid line*; n = 10 per group). Representative sagittal H&E and tdTomato (“tdTom”) immunohistochemistry (IHC) lung sections alongside high powered (100um scalebar) H&E and synaptophysin (Syp) IHC from the LUAD model initiated from AT2 cells on DOX (*above*) or the SCLC model initiated from PNEC cells off DOX (*below*). D) Mean imputed expression of AT2 and PNEC lineage markers (**table S1**) (*90*) in single cells isolated from LUAD (*red;* n = 5,394 cells) or SCLC (*blue;* n = 4,371 cells) ERPMT models with overlaid kernel density estimates reflecting cell density; tdTom+ tumor cells sorted and pooled from n = 3 mice at 8wks post-infection.

LUAD and SCLC tumorigenesis developed with similar penetrance and latency in the ERPMT model. To transcriptionally compare these models, we isolated tdTom+ cells in the on DOX group from the AT2 lineage (labeled hereafter as LUAD; *red*) and the off DOX group from the PNEC lineage (labeled hereafter as SCLC; *blue*) at 8wks on study and performed single cell RNA-sequencing. Expectedly, these models were transcriptionally distinct (**fig. S1A**) and closely resembled their precursors (**Fig. 1D** and **fig. S1, A** to **B**). LUAD exhibited mixed AT2 and AT1 character, consistent with prior observations in the regenerating airway (*22*) and in murine (*12*) and human tumors (*23*). Compared to LUAD, SCLC had lower expression of components of the major histocompatibility (MHC) class II antigen pathway (**fig. S1C**); consistent with MHC class II deficiency in primary SCLC (*24, 25*) and AT2-specific expression of MHC Class II molecules (*26*). Instead, most SCLC tumor cells expressed factors that suppress Notch signaling, including the inhibitory ligand *Dll3* and transcription factor *Hes6* (*27*). Conversely, the LUAD model was enriched for Notch stimulating factors, consistent with this model presenting as a non-neuroendocrine, alveolar-derived LUAD (**fig. S1D**). Additionally, using ATAC-sequencing, regions of chromatin that mapped to *Notch2, Hes1,* and *Sftpc* were differentially accessible in LUAD compared to SCLC (*28, 29*). In contrast, the SCLC model had accessibility in regions controlling neuroendocrine genes, including *Insm1, Chga,* and *Ascl1* (**fig. S1E**). Furthermore, differentially accessible peaks in the MAPK-driven LUAD were significantly enriched for AP-1 motifs like *Jun* and *Fos* whereas several bHLH regulatory elements involved in neurogenesis (*Tcf21*, *Tcf4,* and *Ascl1)* enriched in the SCLC model (**fig. S1F**) (*30*). Taken together, we generated two histologically distinct lung cancers that are united by the process of HT – prompting us to interrogate whether the LUAD model could be encouraged to transform to SCLC.

### EGFR removal and the emergence of neuroendocrine character

We speculated the ERPMT model could be used to understand conversion of LUAD to SCLC, provided adequate selective pressure was applied against the LUAD driver. Following the development of late-stage LUAD, we randomized cohorts to one of three arms: remain on DOX (EPRMT on), come off DOX for 1mo and then re-start DOX (ERPMT on > off > on), or come off DOX and remain off for the duration of the study (ERPMT on > off). Terminal lung cancers developed at statistically different rates across these perturbations (**Fig. 2A**). Critically, if mice had DOX permanently removed, lung tumors were consistent with SCLC, but if DOX was re-started (ERPMT on > off > on), tumors were consistent with LUAD, albeit with fewer, larger lesions (**Fig. 2B**). To better understand the proliferative nature of the residual disease we randomized ERPMT mice developing LUAD to come off or stay on DOX following three daily pulses of the S-phase label EdU. Three weeks later we labeled again, now with a different S-phase analog (BrdU), thereby labeling cells with a proliferative history that continued to cycle following removal of DOX (**fig. S2A**). Macroscopic disease had regressed in groups of mice where DOX was removed (**fig. S2B**). Tumors that remained on DOX were EdU+/BrdU+, whereas off DOX, residual tdTom+ cells were present as single cells and labeled EdU+/BrdU-. This suggested that residual cells did not cycle following DOX removal (**fig. S2, C** to **D**).

**Figure 2.**
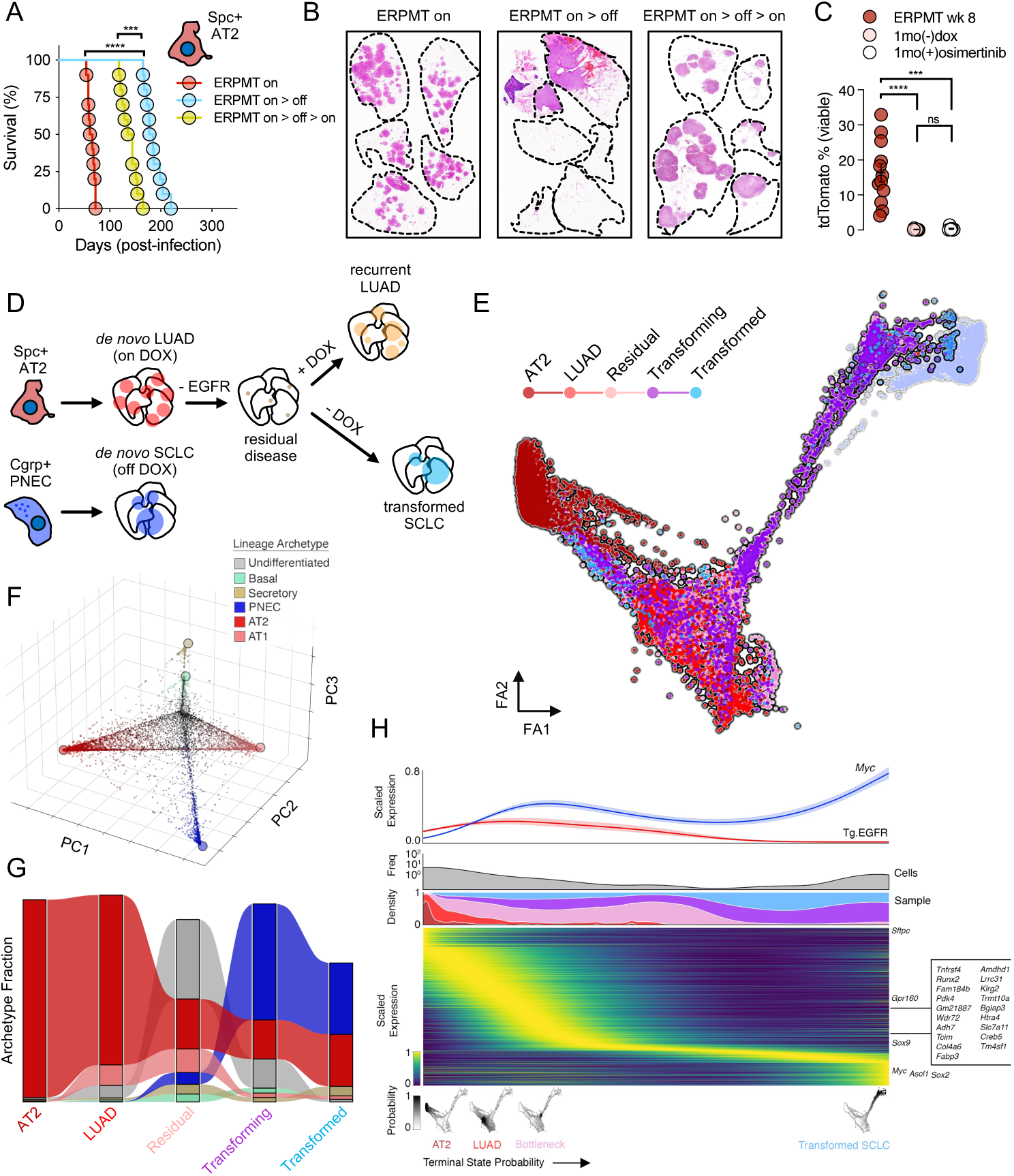
Tracing the origins and transitions between LUAD and SCLC *in vivo*. A) Survival of AT2-derived ERPMT model on DOX (*red*), after DOX removal at 8wks (*teal*), and after restarting DOX diet one month following initial DOX removal (*yellow*; n = 10 mice per group); ***p<0.001, ****p<0.0001. B) Representative sagittal lung H&E sections from moribund mice in each group in *A*. C) tdTom+ burden in the airway after one month of EGFR inhibition via DOX removal (*pink*; n=9) or daily treatment with osimertinib (10mg/kg; PO d1-5 of 7; *white*; n = 9) as compared to 8wks timepoint before DOX removal (*red*; n = 12). DOX removal or osimertinib treatment [on DOX] were initiated at 8wks following the development of extensive ERPMT-derived LUAD tumorigenesis. For clarity, comparisons are effectively an 8wk pre-treatment cohort to ∼12wk post-treatment cohorts (1mo of treatment); ***p<0.001, ****p<0.0001. D) Outline of single cell samples sequenced at distinct timepoints along the transition between LUAD and SCLC following DOX perturbations (*as in A*) E) Force-directed layout of cell states captured along the transition from at AT2 cell to ERPMT LUAD and finally towards a neuroendocrine fate as compared to *de novo* SCLC tumorigenesis (*transparent blue*); colored by sample (*annotated in D*; n = 15,828 cells pooled from 3 mice per sample). F) PCA projection of the lung epithelial lineage probability space (*see Methods*; *same data as in E)*. Individual cells are colored by their max lineage probability and archetypes (*see Methods*) are overlaid as colored nodes (**fig. S4, C** to **E**). G) Flow plot showing relative abundance of cells assigned to their nearest lineage archetype, ordered by sampling time. Bar height is normalized by sample size (log-scale). H) Heatmap of scaled imputed and min-max normalized highly variable transition genes (*see Methods*), top ranked macrostate genes (n = 5, *see Methods*), and lineage markers along terminal cell state probabilities computed using *CellRank* (*see Methods*) for all cells in *E*. For each gene, expression was smoothed along the terminal state probability using a generalized additive model (GAM) as described in *CellRank* (*34*). The top 20 HVGs correlated with the bottleneck macrostate (**table S2**) and select lineage-specific markers are labeled on the right. A stacked KDE reflecting sample abundance (*lower*) and a KDE reflecting total cell frequency (*upper, grey*) are shown above the ranked heatmap. *Above this*, scaled imputed gene expression trends for the oncogenic drivers *Myc* and *EGFR,* modeled using a GAM along the terminal probability.

Transcriptional manipulation of *EGFR* provided a useful genetic tool but did not reflect the clinical scenario where targeted therapies against the EGFR kinase domain are used as the primary therapeutic strategy. To address this, we compared inhibiting the kinase activity of EGFR using osimertinib to the effect of removing DOX, thereby gradually extinguishing *EGFR* expression. After 8wks of LUAD development, the frequency of tdTom+ cells in the airway of ERPMT mice ranged from ∼5-35%. However, this was reduced to <1% in all animals following 1mo of treatment with osimertinib (*31*) or the removal of DOX (∼12wks; **Fig. 2C**). We isolated tdTom+ cells from these groups and found genetic as opposed to pharmacologic suppression of EGFR led to a greater increase in *Ascl1* expressing cells, the definitive lineage marker of the PNEC (*32*) (**fig. S3, A** to **B**). Critically, we did not observe cells expressing both *Ascl1* and *EGFR*, suggesting that a dual-positive state is either short-lived or unviable. Furthermore, independent of *Ascl1* expression, residual tumor cells retained expression of AT2 lineage markers and lacked high expression of a proliferative program characteristic of *de novo* SCLC (**fig. S3C**). These data provided the rationale to explore LUAD to SCLC HT in our ERPMT model by extinguishing oncogenic EGFR through the removal of DOX.

### An undifferentiated, stem-like state emerges during HT

To generate conditions likely to favor HT, we removed DOX from ERPMT mice with late-stage LUAD and transcriptionally profiled single tdTom+ cells from pools of mice (n = 3 per pool) over time (5 timepoints) after *EGFR* removal (**Fig. 2D***; see Methods*). We observed a continuous cell state transition from a normal AT2 cell, through EGFR*-*driven LUAD, and finally towards transformed SCLC at later timepoints. Notably, a minority of tumor cells isolated from samples off DOX formed a bottleneck adjacent to the *de novo* LUAD model prior to breaking out towards the *de novo* SCLC model (**Fig. 2E**). Five extreme phenotypic states (“archetypes”; *see Methods*) (*33*) were identified that corresponded to mature lung epithelial lineages, including AT1, AT2, PNEC, secretory and basal cell types (**Fig. 2F** and **fig. S4B**); however, one archetype exhibited features of a highly undifferentiated state (**Fig. 2F** and **fig. S4C**) and conspicuously mapped to the bottleneck between *de novo* LUAD and transformed SCLC (**Fig. 2, F** to **G** and **fig. S4, B** to **D**).This undifferentiated archetype was not associated with any lung epithelial lineage, but instead expressed modest levels of basal stem cell programs, as well as *Myc* and *Sox2* target genes (**fig. S4C**). Notably, it retained features of the original AT2 lineage, but had yet to express neuroendocrine markers, including *Ascl1* (**fig. S4E**). Tumor cells belonging to this undifferentiated state were a minority in primary LUAD, expanded in the residual LUAD (1mo off DOX), and diminished in the transforming (2mo off DOX) and transformed (>3mo off DOX) populations, which were composed largely of neuroendocrine tumor cells (**Fig. 2G**).

To dynamically model the transition from an AT2 cell to an EGFR-driven LUAD, and finally a transformed neuroendocrine state, we applied *CellRank* (*34*) for single-cell fate mapping. Four stable cellular phenotypes (“macrostates”) were identified along this trajectory capturing the normal AT2, *de novo* LUAD, the HT bottleneck, and finally a transformed SCLC population, respectively (**Fig. 2H***, below heatmap*). MAPK pathway activity (*35*) decreased along this trajectory; conversely, we observed a time-dependent increase in *Myc* transcriptional output (**Fig. 2H***, above heatmap*). Thus, as the LUAD oncogenic driver is removed, a highly specific bottleneck to transformation emerges that is stem-like (*Tm4sf1*) (*36, 37*) and highly proliferative (*38*), with features of neuronal differentiation (*Creb*) (*39–41*) (**Fig. 2H**) and *Myc* downstream signaling (**fig. S4F**).

Transcription factor regulatory modules (“regulons”) that characterize this bottleneck are likewise associated with neuronal plasticity (*Creb5*), airway stemness (*Sox9*) and pulmonary basal cell function (*Trp63*) (**fig. S4G**, *see Methods*). Thus, on the path of HT, as levels of *EGFR* transcript wane, there may be selection for a cell lineage most fit to be driven by high levels of *Myc* (**Fig. 2H** and **fig. S4F**) and cells that break through this bottleneck may be rapidly transformed by *Myc* towards a neuroendocrine fate (**Fig. 2H**).

### AT2 and PNEC lineage driver differences

To directly compare transformation efficiencies between AT2 and PNEC cells by different oncogenic drivers, we infected RPMT mice (which differ from ERPMT mice by lack of the oncogenic *EGFR* transgene) with equal titers of Ad5.Spc-Cre (for AT2) or Ad5.Cgrp-Cre (for PNEC). Initially the tdTom+ frequency was greater in the airway of mice infected with Ad5.Spc-Cre, consistent with a higher baseline frequency of AT2 cells in the lung. However, this was short-lived, followed by exponential expansion of the Ad5.Cgrp-Cre group (**Fig. 3A**). At 8wks post-infection, macroscopic disease was clearly visible in the PNEC-derived RPMT model, but not the comparator (**Fig. 3B**). To simplify cell of origin differences in single oncogenic driver tolerances, we generated mice to temporally trace the effects of oncogenic *Myc*^T58A^ or *EGFR*^L858R^ from the neuroendocrine (*Ascl1*^CreERT2^) or AT2 (*Spc*^CreERT2^) lineages. Following tamoxifen administration to activate Cre^ERT2^, we found that Myc expanded the airway Ascl1+ population and EGFR led to an eventual decline. Conversely, EGFR expanded the AT2 lineage, where Myc was detrimental (**Fig. 3C**). Following survival in a larger cohort of mice, we observed that *Myc* expression from Ascl1+ lineage was sufficient to produce a lethal, fully penetrant phenotype, whereas *EGFR* was not. In contrast, *EGFR* expression alone was sufficient to transform the AT2 lineage, but *Myc* was not (**Fig. 3D**). These data strongly support cell lineage-specific differences in oncogenic driver tolerance of *Myc* and *EGFR* in the lung.

**Figure 3.**
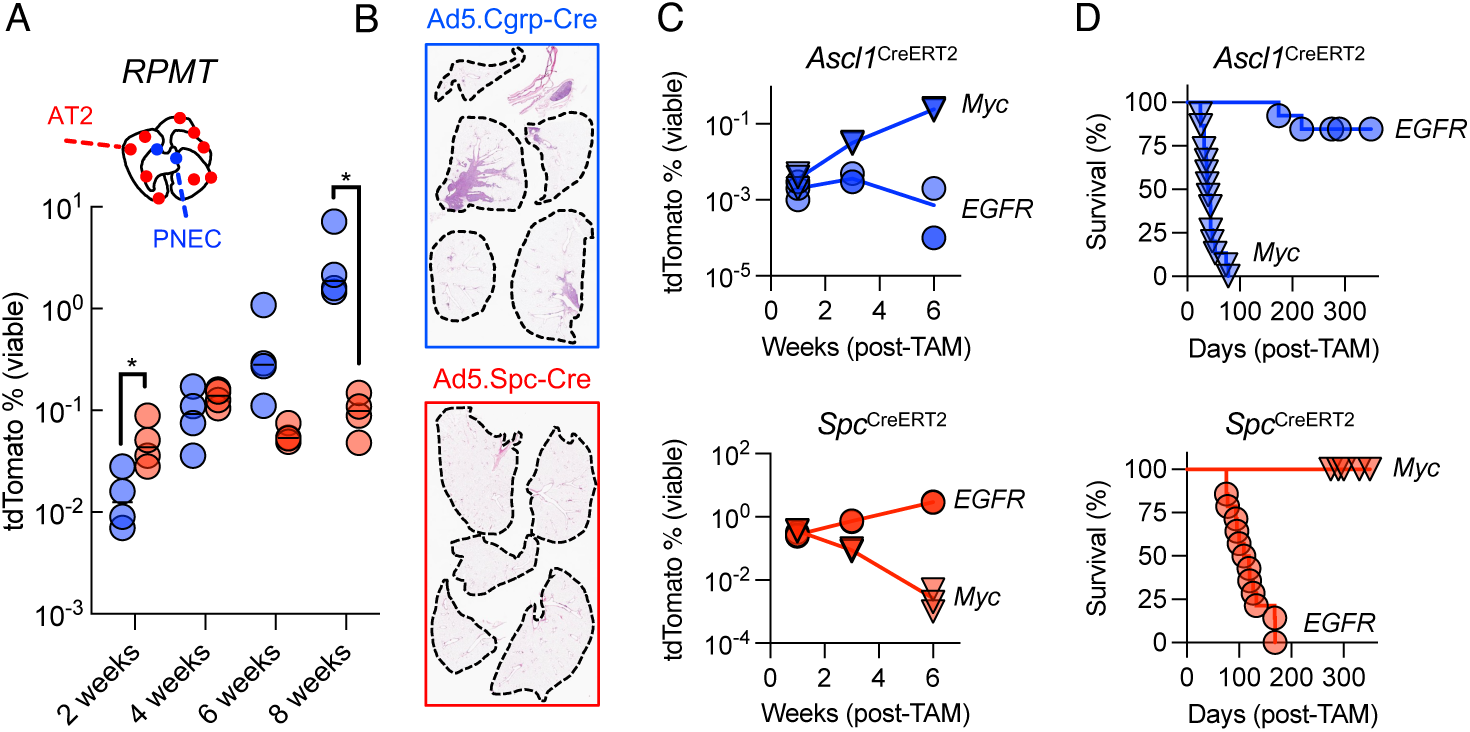
Cell of origin and oncogenic driver incompatibility. A) Frequency of tdTom+ cells in the airway of RPMT mice following infection with equivalent titers of adenovirus (∼10^6^ pfu per mouse) using Ad5.Cgrp-Cre (*blue*) or Ad5.Spc-Cre (*red*) over a period of 8 weeks; n = 4 mice per timepoint); *p<0.01. B) Comparative histology of RPMT (no *Tg.TetO- EGFR^L858R^*allele) mice 8wks post-infection following infection with Ad5.Cgrp-Cre (*blue outline*) or Ad5.Spc-Cre (*red outline*). C) Lineage tracing oncogenic Myc^T58A^ (*down-pointing triangles*) or EGFR^L858R^ (*circles*) on AT2 (*Spc^CreERT2^*; *red*) or PNEC (*Ascl1*^CreERT2^; *blue*) cells in the airway over time; n = 3 mice per timepoint. D) Long-term survival for cohorts shown in *B*; *Ascl1*^CreERT2^ > EGFR^L858R^ (n = 13) or Myc^T58A^ (n = 15) and *below, Spc*^CreERT2^ > EGFR^L858R^ (n = 14) or Myc^T58A^ (n = 12). Mice having a single copy of *Rosa26*^LSL-tdTom^ and *Rosa26*^LSL-rtTA3^ were maintained on DOX chow throughout studies investigating lineage trace allele-mediated expression of *Tg.TetO-EGFR^L858R^*.

The phrase “terminal neuroendocrine” has been used to describe the eventual histology that the ERPMT model transitions towards following *EGFR* withdrawal; however, it was unclear whether this was a terminal state, or if we could study HT in reverse – converting a SCLC tumor towards a LUAD state. Like earlier experiments, we initiated tumorigenesis from PNECs in the ERPMT model and followed cohorts of mice that were on or off DOX from the time of infection. There was a significant delay in the lethality of the model on DOX (**fig. S5A**); however, in lungs of mice on DOX where EGFR protein should be produced, we observed low to absent EGFR by immunohistochemistry (IHC) in tdTom+ tumor regions, morphologically consistent with SCLC (**fig. S5B**). Immunofluorescence for EGFR^L858R^ and Ascl1 showed patchy regions of EGFR-positivity that were excluded from larger areas of Ascl1-positivity (**fig. S5C**), with the Ascl1-positive/EGFR-negative SCLC component of these tumors having the greater burden in all mice examined. Taken together, these data support the conclusions that cells in the PNEC lineage resist transformation towards an EGFR-driven LUAD state, just as cells in the AT2 lineage cannot be easily transformed to SCLC, even though the latter form of HT can occasionally occur under certain conditions. We argue that these intolerances are rooted in incompatibilities between oncogenic drivers and lineage-specific traits.

Oncogenic EGFR led to a gradual elimination of PNECs over months, without evidence of acute intoxication (**Fig. 3C**). It was unclear whether elevated signaling through the MAPK pathway (via EGFR) was a disfavored situation for the PNEC, or if EGFR incompatibility arose through some other mechanism. If excessive MAPK signaling suppressed PNEC proliferation, then inhibition of this pathway should increase Ascl1+ cells. We traced Ascl1+ cells and randomized mice to receive a diet formulated with or without the MEK inhibitor (MEKi) trametinib (**fig. S6A**). We terminated the study after 3mo on MEKi due to toxicity, with adult mice experiencing weight loss approaching our protocol limits (**fig. S6B**). However, no significant differences were observed in the abundance of tdTom+ cells between the two cohorts (**fig. S6C**). Moreover, we did not observe any change in the location or proliferative status of tdTom+ cells in mice treated with MEKi diet (**fig. S6D**), suggesting that physiologic Mek>Erk signaling was not suppressing proliferation of PNECs.

### Myc is sufficient to transform the PNEC

Lineage tracing demonstrated a clear difference in the sensitivity of the AT2 and PNEC cell types to oncogenic transformation by Myc; however, a hallmark of SCLC is inactivation of the *RB1* and *TP53* tumor suppressors, in addition to heightened expression of a Myc family member and its transcriptional targets (*7*). Inspecting the bronchioles of *Ascl1*>Myc mice 1mo post-labeling revealed clusters of cycling, tdTom+ cells that had not yet invaded surrounding tissues, consistent with carcinoma *in situ* (**fig. S7A**). Instead, most *Ascl1*>Myc mice were dying of cancer localized to the thyroid (**fig. S7B**). To date, we have been unsuccessful in activating the *Ascl1*^CreERT2^ allele specifically in the lungs, while sparing the trachea and thyroid. We therefore isolated tdTom+ cells from the airways of four distinct genotypes of mice combining Ascl1 traced *Myc* induction with loss of *Rb1*, *Trp53* or both tumor suppressor genes. We expanded cells *ex vivo* using organotypic culture conditions and engrafted equivalent cell numbers into the flanks of athymic mice. Combined loss of *Rb1* and *Trp53* accelerated the growth of these tumors, but all genotypes were sufficient to form transplantable cancers. Moreover, all tumors had a similar histologic appearance consistent with high-grade neuroendocrine cancer (**fig. S7C**). These data suggest that PNECs could be transformed by Myc [alone] if expanded *ex vivo*, but this did not yet demonstrate tumor maintenance was dependent on Myc.

To test whether PNEC-derived tumors are dependent on Myc, we sorted tdTom+, rtTA3-expressing PNECs from the airway and infected cells with lentiviruses containing tetracycline-promoter driven Myc constructs. Coupling rtTA3 expression to the lineage trace removed the likelihood of infecting lineage-negative cells present as contaminants in cell sorting. While these cells are sparse (**fig. S8**), we could consistently generate organoid cultures from as few as 50 cells when DOX was present in the culture media to transcriptionally drive *Myc*. Removal of DOX resulted in near complete growth suppression of organoids expressing Myc^WT^ as compared to Myc^T58A^ (**Fig. 4, A** to **C**), a long-lived version of Myc resistant to GSK3b- mediated phosphorylation and eventual degradation (*42*) (**Fig. 4C**). Engrafting PNECs expressing inducible Myc^WT^ into immunocompromised mice demonstrated DOX-dependent tumor growth. Removing DOX from tumor-bearing mice dramatically reduced tumor volumes over the course of several weeks (**Fig. 4D**), with residual, fibrotic tumor tissue mostly comprised of non-cycling cells (**Fig. 4E**). Consistent with their rapid proliferation, Myc-driven PNEC organoids were sensitive to compounds that exacerbated replication stress such as topoisomerase inhibitors (etoposide), as well as inhibitors of enzymes required for cell cycle progression, including Cdk4/6 (palbociclib) and Wee1 (adavosertib) (**Fig. 4F**). Additionally, direct inhibition of the Myc-Max interface using the small molecule MYCi975 (*43*) provided similar reduction in organoid growth as compared to DOX removal. (**Fig. 4F**).

**Figure 4.**
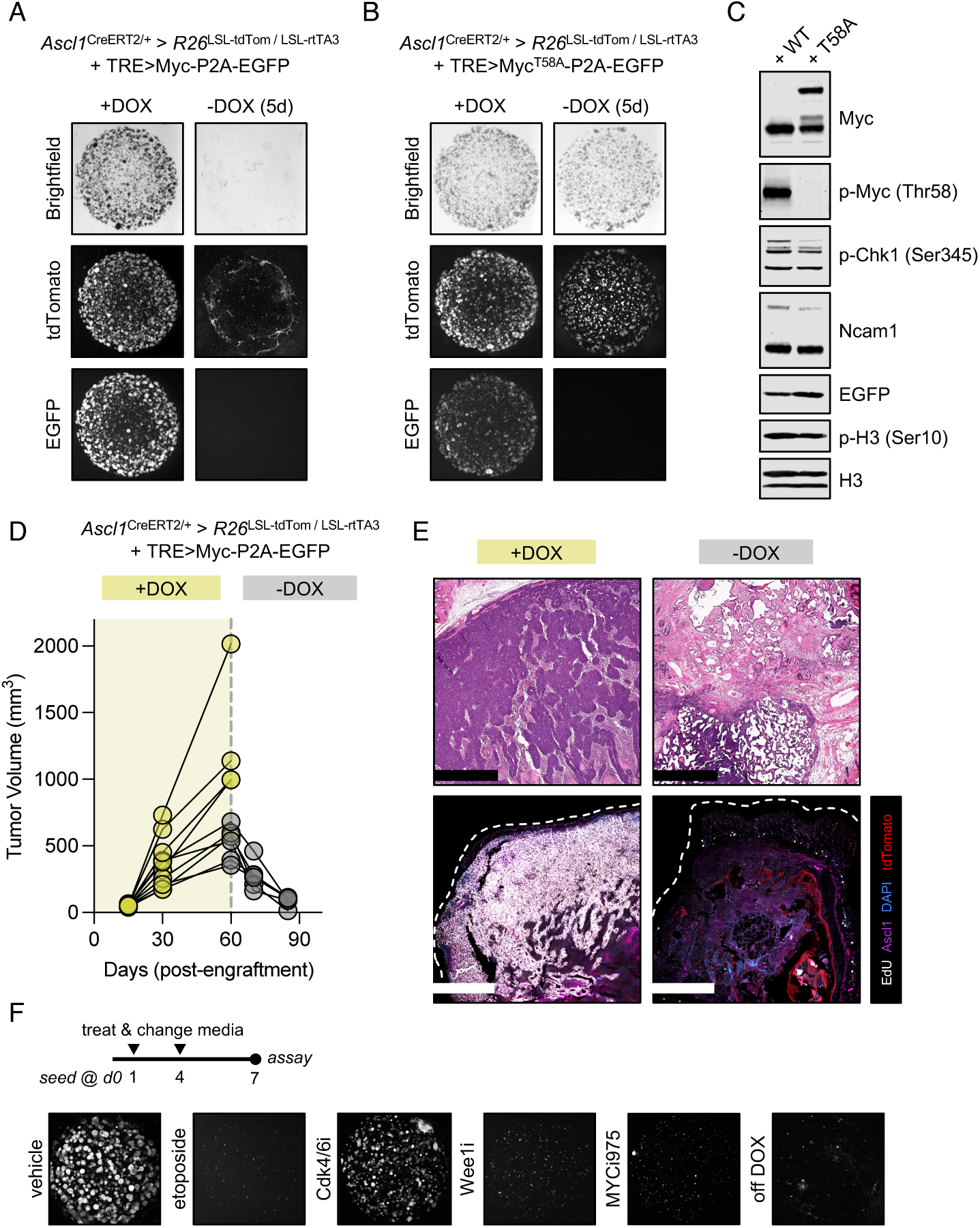
Proliferation of the pulmonary *Ascl1*+ lineage is Myc-dependent and sufficient for tumorigenesis. A) Lineage labeled Ascl1+ cells from the lung expressing tdTom and rtTA3 were sorted then infected *ex vivo* with lentiviral constructs expressing a *Myc* gene tied to *EGFP* under control of a tetracycline-regulated promoter. Appearance of spheroid cultures +/- DOX at 5d of culture; single 50uL spheroid area shown. B) Appearance of spheroid cultures +/- DOX at 5d of culture from Myc^T58A^ construct; single 50uL spheroid area shown. C) Western Blot for protein markers of proliferation, neuroendocrine identity, cellular stress and Myc phosphorylation; organoid cultures were collected 5d after seeding 10K cells in 50uL droplets of 50% Matrigel® and cultured in the presence of 1ug/mL DOX, where media was changed every other day. D) Engraftment of cells from *A* into immunocompromised mice (athymic/nude) on DOX diet show *in vivo* growth dependent on DOX diet; dashed line indicates point of DOX chow removal in ∼50% of cohort (n = 5 per arm). E) Gross appearance (H&E; 1mm scalebar) and immunofluorescence for lineage and proliferation markers in tumors on DOX at volumetric endpoints (∼1000mm^3^) or off DOX at point of maximal tumor consolidation (2wks off DOX; 1mm scalebar). F) Experimental timeline for *Ascl1*>TRE>Myc;tdTom organoids from *A* treated with indicated compounds (DOX-containing media) every 3d for one week before imaging representative wells at day 7; cells seeded in media without DOX included as negative control.

### AT2 cells are refractory to transformation by Myc

While Myc alone may be sufficient to drive transformation and expansion of PNECs, our earlier results with lineage tracing suggested it was insufficient to transform the AT2 lineage (**Fig. 3C**). To further investigate what underlies this bottleneck, we established organoid cultures using recently published methods (*44*) from sorted, normal AT2 cells or those expressing *Myc^T58A^*. AT2 cells expressing *Myc^T58A^*expanded rapidly as compared to their wildtype counterparts but were unsustainable beyond 3 passages (**fig. S9A**). In early timepoints following the expression of *Myc^T58A^* in AT2 cells *in vivo*, we noted incorporation of EdU in tdTom+ cells, but at a year following the initiation of the trace, remaining tdTom+ cells in the airway failed to incorporate EdU, suggested they were no longer proliferative or had been eliminated (**fig. S9B**). *Ex vivo*, AT2 organoids expressing *Myc^T58A^* demonstrated increased DNA damage sensing, replication stress and markers of programmed cell death as compared to wildtype (**fig. S9C**). It is unlikely this represented an excess of Myc protein, as levels were significantly lower in AT2 cells compared to levels tolerated in PNEC organoids (*45*) (**fig. S9D**). Finally, consistent with the observation that Ras signaling through PI(3)K relieves Myc-induced apoptosis (*46*), we likewise observed that Myc significantly accelerated oncogenic EGFR-driven LUAD (**fig. S9, E** to **F**).

While our data suggest that oncogenic drivers of the AT2 and PNEC lineages are in stark contrast, it remained unclear whether other epithelial airway cells can transform in response to *Myc* alone. To address this, we performed a generalized, conditional trace using an *Nkx2.1*^CreERT2^ allele that will drive Cre-mediated recombination in a broad range of cell types derived from the anterior foregut endoderm, including the trachea, pituitary, thyroid and most of the lung (*47*). At early timepoints, lineage-labeled tissues within the thyroid (both Ascl1+ and Ascl1-) expanded following *Myc* expression compared to control mice (**fig. S10A**); however, at later timepoints there was outgrowth of tdTom+/Ascl1+ cells in bronchioles not observed in wildtype controls (**fig. S10B**). Together these data suggest the Ascl1+ PNEC is unique in its tolerance to Myc, but we had not yet explained how intolerance to Myc could be overcome in the AT2 lineage during HT.

### Pten deletion removes a barrier to Myc transformation

Genes upregulated in tumor cells as they escape the bottleneck to HT in our ERPMT model (termed “*breakout*”; **fig. S11A**) were notably associated with PI(3)K signaling (**fig. S11, B** to **C**) as compared to cells found in the bottleneck and unable to adopt neuroendocrine fate (*48*). Thus, we asked whether increasing PI(3)K-dependent Akt signaling through deletion of *Pten* would enable *Myc^T58A^*-driven transformation in an AT2 lineage trace. Strikingly, we observed fully penetrant *Myc*-driven transformation in an AT2 cell when *Pten* was inactivated (**Fig. 5A**). At the median period where animals in the *Spc*>Pten;Myc cohort were moribund, lungs from *Spc*>Myc mice showed no evidence of macroscopic disease (**Fig. 5B**). Similar observations were made combining *Myc* expression with a mutant, actived PI(3)K allele (E545K) or inactivating both copies of *Pten* (**Fig. 5B**). Examining *Spc*>Pten;Myc animals 3mo post-recombination reinforced the observation that lesions were variable in their frequency, size and histologic appearance (**fig. S12**). However, they were Ascl1-negative with glandular appearance, suggesting they were adenocarcinoma-like and excluding the likelihood of SCLC, squamous or mixed histology (**fig. S12**). AT2 lineage markers, including pro-surfactant protein C (pro-SPC) and MHC Class II, were low or absent and basal epithelial markers like Sox2 and keratin 18 were variably expressed. We also observed a notable increase in the highly undifferentiated, basal-stem like state following combined loss of *Pten* and expression of *Myc* in AT2 cells *–* not seen with either perturbation alone (**Fig. 5C**). As expected in response to the loss of *Pten*, these basal-like cells exhibited higher levels of Myc target genes, PI(3)K signaling and stemness signatures (**Fig. 5D** and **fig. S13A**) following cellular adaptation to *Myc*.

**Figure 5.**
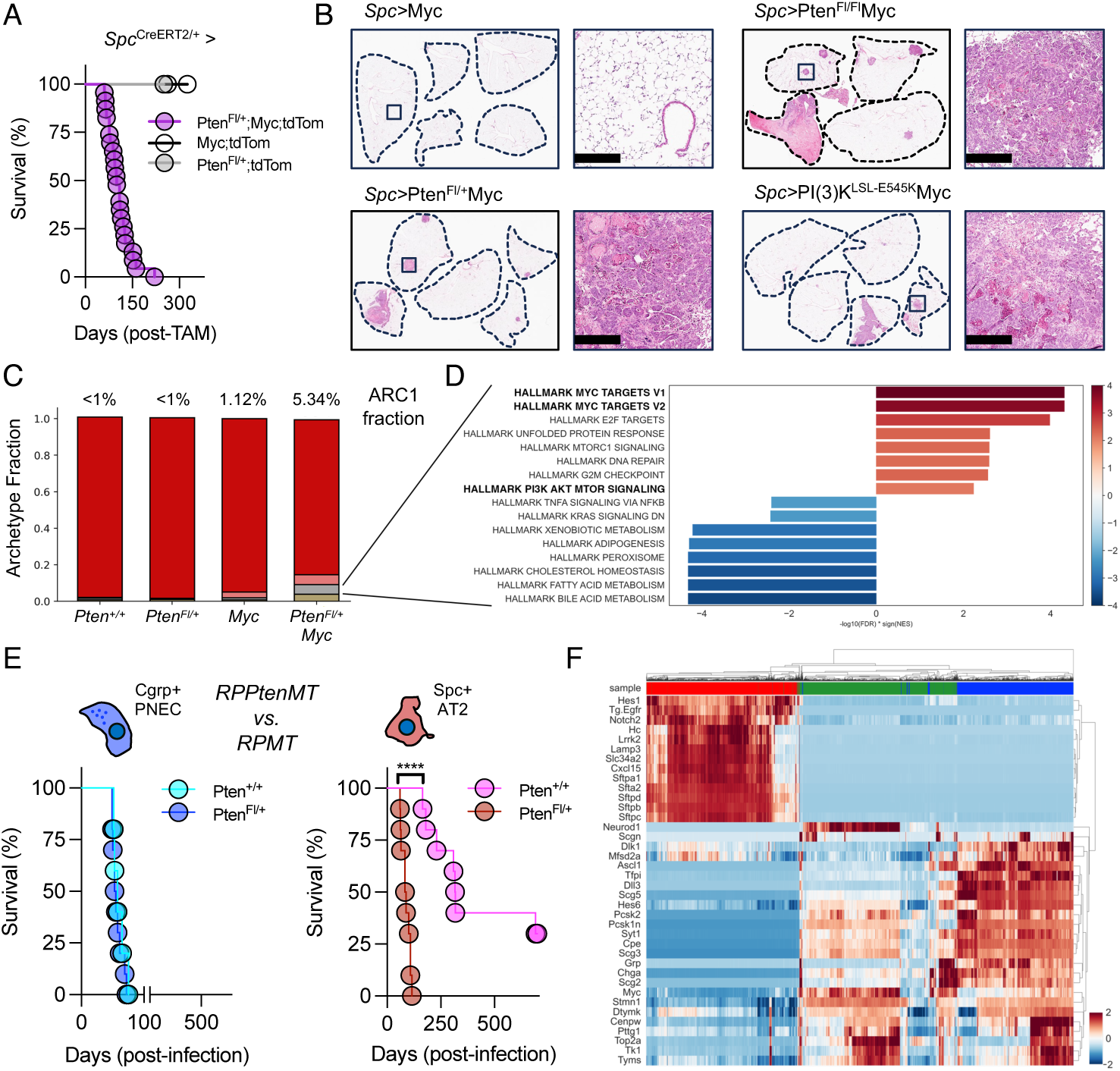
Loss of *Pten* in the AT2 lineage removes the barrier to Myc transformation. A) Survival of mice where Myc (n = 12), Pten^Fl/wt^ (n = 5) or the combination of these alleles (n = 23) are initiated in AT2 cells using a single copy of *Spc*^CreERT2^. Data were censored between 250-325d in the non-lethal arms. B) Comparative histology of whole lungs from representative *Spc*>Myc, *Spc*>Pten^Fl/+^;Myc), *Spc*>Pten^Fl/Fl^;Myc or *Spc*>PI(3)K^LSL-E545K^;Myc mice at ∼3mo post-labeling. Higher magnification regions (*boxed*) are provided at *right*; 200um scalebar. C) Bar plot showing fraction of each epithelial lineage archetype detected per sample as in (**Fig. 2G**). D) Bar plot of Hallmark gene sets significantly enriched (FDR < 0.01) within the undifferentiated cell state of the *Spc*>Pten^Fl/+^;Myc sample pool, as in (**fig. S4F**). E) Effect of *Pten* deletion combined with deletion of *p53*, *Rb1*, and expression of *Myc* and *tdTomato* (RPPtenMT or RPMT) in neuroendocrine (*blue*) or AT2 (*red*) cells; n = 10 per arm with x-axis split for clarity, ***p<0.001. F) Clustered heatmap of imputed and z- normalized expression of AT2 and PNEC signature genes, model oncogenic drivers (*Myc* and *Tg.EGFR*), and SCLC subtype identifiers (*NeuroD1*, *Pou2f3*, and *Yap1*) for all tumor-epithelial cells from the RPPtenMT model (*green*) and the *de novo* LUAD (*red*) and SCLC (*blue*) models described (**Fig. 1**). Genes and cells are clustered using the Euclidean distance method.

### Pulmonary basal cells efficiently generate SCLC

The basal stem-like transcriptional program associated with AT2 cells capable of adaptation to *Myc* supported the possibility that an intermediate state during HT may be basal-like. More generally, this raised the possibility that the basal cell may serve as a cell of origin for SCLC, as speculated by others (*49*); However, targeting basal cells in mouse models is difficult due to their anatomic location, noted to be refractory to viral infections delivering Cre (*50*). Indeed, we observed an absence of tdTom+ cells in the lungs of mice one-month post-labeling when using a *Krt5*^CreERT2^ allele to target basal cells (**fig. S13A**). However, following regeneration in the proximal lung induced by naphthalene damage of secretory cells (*51, 52*), tdTom+ cells were detected within the lungs of mice, without a significant change in the trachea or thymus (**fig. S13B**). As basal cells are known to serve as multi-potent progenitors (**fig. S13C**) (*51–53*), we then crossed mice to lose *Rb1* and/or *Trp53* from the basal lineage trace and found *Rb1* loss alone was sufficient to skew cells towards a neuroendocrine fate (**fig. S13D**).

Consistent with this, we observed fully penetrant neuroendocrine tumorigenesis in mice that have lost both *Rb1* and *Trp53* with (*Krt5*>RPMT) or without (*Krt5*>RPT) expression of Myc from the basal lineage (**fig. S13E**); however, tumors arose with significantly shorter latency in mice producing oncogenic *Myc*. Expression of the conditional reporter allele was easily observed throughout the keratinized epithelium of mice following labeling, but interestingly, tumorigenesis was restricted to the proximal airway (**fig. S13, F** and **H**).

Histologically, both models produced SCLC; however, we noted that most tumors were adjacent to or involving the thymus and did not invade the lungs, unless we damaged lungs with naphthalene (**fig. S13, G** to **I**). These data suggest basal cells can not only serve as cells of origin for SCLC, but basal progeny and anatomic location can be influenced by genotype and injury.

### Rb1 loss is required for LUAD to SCLC HT

Deletion of *Pten* was sufficient to lower the barrier to transformation by Myc in the AT2 lineage; however, the resulting tumors were not neuroendocrine (**fig. S12**). We suspected the loss of *Rb1* was required (*10*) – in addition to the suppression of *EGFR* and adaptation to Myc – for the emergence of neuroendocrine disease. To test this, we generated a model in which *Rb1* was wildtype (“EPMT”; *Rb1*^+/+^) and initiated tumorigenesis in AT2 cells using Ad5.Spc-Cre or a lineage trace allele (*Spc*^CreERT2^>EPMT). On DOX, both models produced aggressive LUAD with the difference in latency likely reflecting the broad initiation achievable using a lineage trace allele compared to sparse infection. (**fig. S14, A** to **B**). Critically, LUAD did not recur in EPMT mice taken off DOX and the residual tdTom+ cells did not express *Ascl1* (**fig. S14, A** to **C**), further suggesting that it is the combined loss of *Rb1* and *EGFR* suppression that are together required for an Ascl1+ neuroendocrine tumor to emerge by HT.

Thus, to recapitulate bona fide transformation from an AT2 cell to SCLC efficiently, we generated a final model in which we could delete *Rb1*, *Trp53*, a copy of *Pten,* and express *Myc* and *tdTom* (abbreviated *RPPtenMT*). If initiated from PNECs there was no difference in latency between the RPMT and RPPtenMT models, but a significant difference emerged if the model was initiated from AT2 cells (**Fig. 5E**). Tumor cells from the AT2-derived RPPtenMT model had expression profiles more like PNECs and clustered near *de novo* SCLC, distinctly away from *de novo* LUAD (**Fig. 5F** and **fig. S15A**). We also noted two subpopulations of cells stratified by expression of *Ascl1* (**fig. S15B**). Differential gene expression in the *Ascl1^low^*group was notable for expression of neuronal genes including, *Stnm2*, *Nfix* and *Mapt*. The *Ascl1^low^* group also expressed high levels of *NeuroD1* – consistent with prior work showing *Myc* driving subtype plasticity from an Ascl1^high^ to Ascl1^low^ / NeuroD1^high^ transcriptional profile in multiple models of SCLC (*54–58*) (**fig. S15C**). Pathology review revealed extensive heterogeneity with mixing of classic SCLC and large cell neuroendocrine (LCNEC) tumor features, marked by variable expression of neuronal and neuroendocrine markers (**fig. S15D**).

Taken together, we conclude that the AT2 cell is highly refractory to transformation by oncogenic Myc, as are many other cell types within the lung – the noted exception being the PNEC. The intolerance of AT2 cells to Myc can be relieved through activation of the Akt signaling pathway, such as the deletion of the tumor suppressor *Pten*, to generate a permissive, stem-like state, and the full conversion to a Myc-driven, high grade neuroendocrine cancer requires the additional loss of *Rb1*.

## Discussion

Carcinogenesis and related processes, like tumor progression, therapy resistance and HT remain incompletely understood. To mechanistically understand HT in lung cancer, we have developed new models of lung tumorigenesis initiated in different pulmonary cells of origin to follow the transformation of EGFR-driven LUAD to Myc-driven SCLC. In doing so, we demonstrate that HT can be simplified conceptually to a change in oncogenic drivers as the tumor cells transition between states that resemble cells in the AT2 and the PNEC lineages. The driver oncogene in the PNEC lineage is Myc and the bottleneck in HT can be relieved by mechanisms that allow an AT2 cell to become a PNEC-like cell that is driven to an oncogenic state by high levels of Myc.

Understanding events that facilitate HT has been limited by the lack of samples obtained from the same patient before, during and after HT. Comparisons of *de novo* SCLC to *transformed* SCLC have highlighted pathways activated following neuroendocrine differentiation (*59*). A more recent study relied on “putative” HT samples where a LUAD oncogenic driver was present in a histologically confirmed SCLC. Interestingly, when compared to *de novo* SCLC, these cases were significantly enriched for mutations that activate the PI(3)K signaling pathway (*60*). Consistent with this, we found that loss of *Pten* or activation of *Pik3ca* were sufficient to break a barrier of the AT2 lineage to transformation by Myc, but insufficient to lead to neuroendocrine transformation. Prior work has established a combination of genetic events capable of reprogramming human cell types to neuroendocrine cancers (*61*); however, the intermediate steps in this process remain unclear. We build upon this work by developing intact genetically engineered animals and sampling tumor cells throughout the lifetime of HT. Moreover, we find that while *Rb1* loss is necessary for HT to a neuroendocrine cancer, it is the extinction of the driver oncogene *transcriptionally* that is required prior to the emergence of neuroendocrine features. These results may provide a possible explanation for earlier work in *Kra*s-driven LUAD where *Rb1* and *Trp53* were inactivated (*62*). While the authors found an increase in tumor histologic heterogeneity following *Rb1* loss, neuroendocrine histology was notably absent. We would suggest this was due to the continued presence of the LUAD driver.

Beyond the genetic manipulations and changes in gene expression described here, we speculate that there are likely other mechanisms that allow for such lineage conversion, plasticity or trans-differentiation. Lineage tracing has demonstrated that basal progenitors can give rise to PNECs (*63, 64*), and more recently, tracheal basal cells differentiate towards a PNEC fate under hypoxic conditions, but the intermediate transcriptional steps in this complex process are uncharted (*65*). In our attempts to describe the transition state between LUAD and SCLC, we found that these tumor cells appear “basal-like”, based on their transcriptional profile, but lacked definitive basal lineage markers. Instead, these cells are more accurately described as a “lineage negative,” stem-like progenitors. Such cells have been described to arise in the mouse lung following major airway injury (*66*). Thus, the airway cell most capable of such plasticity may be the pulmonary basal cell or a stem-like cell that has yet to be fully described.

Finally, it is unclear whether therapeutic targeting of pathways facilitating HT (such as the Akt pathway) in patients poised to undergo HT will have efficacy. Our results suggest that non-invasive monitoring for activation of the Akt signaling pathway [such as the appearance of *PI(3)K* mutations in circulating tumor DNA] may serve to alert physicians to the likelihood of HT, prior to the emergence of an aggressive, transformed SCLC. It is not yet known whether direct targeting of Myc in such Myc-driven scenarios (including HT or *de novo* SCLC tumorigenesis) will be fruitful; however, pre-clinical Myc inhibitors have advanced substantially over the last decade and might ultimately have clinical utility in multiple contexts (*43, 67, 68*).

## Limitations of the study

There are several key limitations to this study: First, we are attempting to understand HT by modeling a human phenomenon in the mouse where the complexity of cell types and microenvironments are not identical. Recent descriptions of the human distal airway have found a diversity of cell types in human lungs that are not observed in the mouse, including specialized regenerative cells with hybrid alveolar-secretory character and multiple subtypes of basal cells (*19, 69, 70*). For example, basal cells are not present in the mouse lung, unless the airway is damaged, which we demonstrate using naphthalene. Furthermore, most laboratory mice are housed under pathogen-free conditions. We know that lifetime exposure to various carcinogens and particulates fundamentally alters the microenvironment of the lung, including the likelihood that a cancer will develop (*71, 72*). Second, there are processes in human cells not found in the mouse that may be critical for HT, including the role for APOBEC-mediated hypermutation. Studies of EGFR-mutated LUAD have found APOBEC mutagenesis signatures following treatment with EGFR-targeted therapies (*11, 73, 74*) and this mutational signature is enriched post-HT (*1, 60*). It is unclear whether APOBEC mutations are directly responsible for activating the PI(3)K signaling pathway, thereby [hypothetically] relieving intolerance of the AT2 cell to Myc (*1, 59, 60*); however, other studies would suggest this is possible (*75, 76*). Finally, we were unable to initiate tumorigenesis in the ERPMT model using the *Spc*^CreERT2^ lineage trace allele. This proved impossible because, in the mouse genome, the *Rb1* locus is ∼3Mb from the *Sftpc* locus (GRCm38/mm10 build). Such proximity precludes generation of the desired genotype with both homozygous floxed copies of *Rb1* and the *Spc*^CreERT2^ knock-in allele. We address this issue through a combination of genetic “leave-one-out” experiments in tumorigenesis models complemented with single oncogenic driver lineage tracing.

## Funding

National Cancer Institute of the National Institutes of Health grant 5U01CA224326 (to H.V.)

National Cancer Institute of the National Institutes of Health grants P01CA120964 and R35CA197588 (to L.C.C.)

National Cancer Institute of the National Institutes of Health grants R01CA256188-01, R01CA280414-01 and R21CA266660-01 (to A.M.L.)

The Burroughs Wellcome Fund (to A.M.L.)

The Lung Cancer Research Foundation (to A.M.L.)

The Damon Runyon Cancer Research Foundation fellowship DRG-2343-18 (to E.E.G.)

H.V. is the Lewis Thomas University Professor at Cornell University.

## Author Contributions

Conceptualization: E.E.G., H.V.

Methodology: E.E.G., E.M.E., M.J.H., K.L., J.T., B.D.S., L.C.C., A.M.L.

Experimental Design: E.E.G., E.M.E., A.M.L., H.V.

Visualization: E.E.G., E.M.E., M.J.H., A.M.L.

Funding acquisition: E.E.G., L.C.C., A.M.L., H.V. Supervision: H.V., A.M.L.

Pathology review: C.Z. Writing – original draft: E.E.G.

Writing – review & editing: all authors

## Data and Materials Availability

### Lead contact

Requests for resources should be directed to and will be fulfilled by Eric E. Gardner or Ashley M. Laughney.

### Materials

All mouse models, organoids derived from mice and plasmids described in this work will be made available to investigators through an institutional or third-party Material Transfer Agreement (MTA), upon reasonable request. Select plasmids will be submitted to Addgene upon manuscript acceptance. Not all mouse strains are currently active – please contact Eric E. Gardner for specific information and availability.

### Data and code

The processed single cell data and relevant code, including Docker environments with Jupyter notebooks demonstrating key analyses, will be available on the Gene Expression Omnibus (GEO) and Laughney Lab GitHub upon manuscript acceptance.

## List of Supplementary Materials

### Materials and Methods

*Supplementary Documentation Figs. S1 to S15

*Tables S1 to S4

*Notes S1 to S5 References (*77–94*)

*\*not included in pre-print upload*

## Materials and Methods

### Animal Work

#### Genetically Engineered Mouse Models (GEMMs) and Parental Strains

The following strains have been described previously and were available as live colonies at the time of institutional acquisition: *Tg.TetO.EGFR^L858R^* (MGI:3690078), *Rb1*^Flox^ (JAX #008186 & #026563), *Trp53*^Flox^ (JAX #008462), *Pten*^Flox^ (JAX #006440), *Pik3ca*^LoxE545K^ (MGI:5438138), *H11*^LSL-CAG-Myc(T58A)-IRES-Luc^ (JAX #035557 & 029971) *Rosa26*^LSL-tdTomato(Ai14)^ (JAX #007914), Rosa26^CAGs-LSL-rtTA3^ (JAX #029617), *Ascl1*^CreERT2^ (JAX #012882), *Sfptc*^CreERT2^ (JAX #028054), *Krt5*^CreERT2^ (JAX #029155), and *Nkx2.1*^CreERT2^ (JAX #014552) (*20, 77–86*). Our strategy to create the ERPMT model was to first generate mice homozygous for the following [simplified] genotypes “ERPR” (*Tg.TetO.EGFR^L858R^*_;_ *Rb1*^Flox/Flox^_;_ *Trp53*^Flox/Flox^_;_ *R26*^LSL-rtTA3/LSL-rtTA3^) and “RPMT” (*Rb1*^Flox/Flox^_;_ *Trp53*^Flox/Flox^_;_ *H11*^LSL-Myc(T58A)/LSL-Myc(T58A)^; *R26*^LSL-tdTom/LSL-tdTom^) in separate colonies and then cross to use the F1s, if positive for the EGFR transgene. Single alleles were backcrossed against C57BL/6J (JAX #000664). Compound genotypes were maintained on a mixed background containing C57BL/6J, C57BL/6N, FVB/N and 129S. Harem breeders were maintained of a 50:50 mix of standard diet (Purina Pico #5053) and Love Mash rodent reproductive diet (Bio-Serv). Zygosity of lineage trace and reporter alleles are indicated throughout the figures and are in most cases heterozygous or hemizygous. Both male and female adult mice (6-12wks of age when enrolled on study) were included in all experiments. One noted exception was the use of only female, outbred athymic/nude mice in xenograft experiments (Taconic; NCRNU). All animal work, including the use of photography to describe the recombination observed in the *Krt5*^CreERT2^>RPMT model was approved by Weill Cornell Medicine’s Institutional Animal Care & Use Committee (IACUC) protocol #2015-0017. Additional details on cohort size, genotypes, phenotype median latency and experimental details can be found in the ***Supplementary Documentation*** (*see* **table S4**).

#### Diet formulations

Trametinib (LC Labs) diet was formulated at 5mg/kg in AIN-76A base diet (Bio-Serv). Doxycycline diet was formulated by LabDiet® (625ppm). All diets were gamma irradiated by the provider.

#### Mouse lung collection, enzymatic digestion, flow cytometry and cell sorting

In brief, mice were euthanized by inhaled CO2 for 5’ and then quickly sheathed and dissected to expose the chest cavity. 10cc of warm PBS + 1mM EDTA was used to perfuse the ventricle and clear the lungs of circulating blood. The trachea was then exposed and a 21G needle was inserted to either 1) inflate the airway with 2cc of 1.5% low melt agarose (Lonza SeaPlaque™; Tm 65°C) for immediate fixation in cold 4% paraformaldehyde (Sigma; pH adjusted to 7.2) or a digestion cocktail composed of collagenase type IV (Worthington), elastase (Worthington), Liberase™ TM (Roche) and DNAse I (Worthington) in Opti-MEM I reduced serum medium (Thermo Fisher Scientific) supplemented with 10uM Y-27632 (Med Chem Express). Lung lobes were grossly dissected away from the tracheobronchial tree and placed into gentleMACS C tubes, making up the final reaction volume to 5cc using digestion cocktail. Enzymatic digestions were performed for one hour on a Miltenyi gentleMACS octo dissociator at 37C. Single cell suspensions were then passed over a 70um mesh filter and washed 3X using MACS buffer (PBS pH 7.4 + 1mM EDTA + 0.5% BSA). If red blood cells were of concern, samples were incubated with hypotonic ACK lysis buffer for 3’ and then washed with MACS buffer until clear. Cells were sorted for viable, single tdTomato+ events on one of the following 4 instruments at the Weill Cornell Medicine Flow Cytometry Core Facility: BD Aria II, BD Influx, BD Symphony S6 or a Sony MA900. Cells were analyzed on a Thermo Fisher Scientific Attune NxT acoustic cytometer.

#### Tissue fixation, immunohistochemistry (IHC) and immunofluorescence (IF)

Lung tissue inflated with low melt agarose was fixed overnight at 4C on a rocking platform. The next day tissues were washed with MilliQ water 3 times for a total of 15’ and then transferred to 70% ethanol on a rocking platform at 25C. Ethanol was changed several times until it ran clear. Embedding, sectioning, routine histologic stains (e.g. H&E) and automated IHC were performed by Histowiz. The following IHC antibodies were used: anti-tdTomato (Rockland #600-401- 379), anti-synaptophysin (Leica #SYNAP-299-L-CE), anti-EGFR (Abcam #ab237986), anti-Mash1/Ascl1 (Abcam #ab74065), anti-Pgp9.5 (Abcam ab108986), anti-Dll3 (Abcam ab229902), anti-MHC Class II (I-A/I-E) (eBiosciences #14-5321-82), anti-keratin 18 (Bioss #BS-5405R), anti-Sox2 (Cell Signaling #149962) and pro-surfactant protein C (Abcam #ab90716). Slides for IF were heated to 55C for 30’ and then de-waxed with two incubations in HistoClear (Leica) and progressive re-hydration from 100% ethanol to 50% ethanol over 20’.

The antigens were unmasked in a pressure cooker set to ramp to 110C for 10’ in an alkaline retrieval buffer (10mM TBS pH 9.0, 0.05% Tween-20 and 1mM EDTA). Following retrieval, slides were cooled for 10’ in a circulating water bath and then washed 1X with TBST (10mM Tris pH 7.2 and 0.05% Tween-20) and blocked for 30’ using a buffer composed of 0.5% BSA, 0.05% Triton X-100 in 1X PBS pH 7.2 supplemented with mouse-on-mouse blocking reagent (Vector Biolabs; 1:50) and Fc receptor blocker reagent (Innovex Biosciences; 1:25). Slides were then briefly washed in TBST and primary antibodies were incubated overnight in blocking buffer at 4C in a humidified chamber. The following IF antibodies were used at the indicated dilutions: anti-Mash1/Ascl1 (Abcam #ab211327; 1:100), anti-RFP (Rockland #600-401-379 or #200-301-379; 1:300), anti-Cyp2f2-A488 (Santa Cruz; #sc-374540; 1:500), anti-BrdU (eBioscience #14-5071-82; 1:500), anti- EGFR^L858R^-PE (Cell Signaling #64716; 1:500), anti-EpCam-A488 (Abcam # ab237384; 1:500). EdU was detected using a Click-iT Plus EdU imaging kit (Thermo Fisher Scientific). Secondary fluorophore-conjugated antibodies were used at a 1:1000 dilution (AlexaFluor dyes; “AF”) and during the secondary antibody incubations (60’ at RT in dark) DAPI was included as a nuclear counterstain (1ug/mL; Sigma Aldrich). Directly conjugated primary antibodies were incubated as a final step prior to washing and mounting (Thermo Fisher Scientific; ProLong Diamond Anti-Fade). Slides were cured overnight in the dark prior to scanning using an Axioscan 7 instrument. Images were analyzed in Zen Connect (Carl Zeiss).

### Primary Cell Culture

#### Organoid culture of AT2 and PNEC cultures

All organoids were seeded in a 50% GFR, phenol ref-free Matrigel (Corning) solution as 50uL droplets and solidified for 10’ in a 37C incubator prior to the addition of media. For the first 2d following seeding and 1d prior to splitting, 10uM Rock inhibitor was included in culture media. Media was changed twice a week and organoids were split when nearing confluence. AT2 organoid cultures were developed as described (*44*), with a few modifications. We preferred the use of acutely traced Spc^CreERT2^ > Rosa26^LSL-tdTom^ cells over indirect immunostaining for MHC class II. The use of Noggin was replaced with 10nM LDN-193189 (Selleck Chem) and A-83-01 was omitted from all culture conditions. PNEC organoids were cultured in modified, serum-free MTEC media supplementing Advanced DMEM-F12 (Thermo Fisher Scientific) with 1X BEGM singlequots™ supplement pack (Lonza; #CC-4175), glutaMAX, penicillin/streptomycin and HEPES (Thermo Fisher Scientific). Cells were isolated from Matrigel through sequential incubations in TrypLE (Thermo Fisher Scientific) at 37C and quenching with PBS. Following the initial wash and spin (500*g for 3’) spent Matrigel was carefully removed by a pipetted and the remaining cell pellet was re-suspended in TrypLE for the second wash. All liberated organoids were passed through 70um mesh filters prior to re-seeding, cryo-preservation or collection for further analyses.

### Molecular Biology

#### Lentiviral constructs and ex vivo infections

Inducible Myc constructs were created from synthetic fragments (Twist Biosciences) sub-cloned into a general backbone derived from LT3GEPIR (gift from J. Zuber; Addgene plasmid #111177) using Gibson Assembly reagents (NEB). Select elements were removed from the backbone (such as rtTA3, PuroR, etc.) using PCR. Crude lentiviral supernatants were generated in HEK-293Ts cells, packaging with psPAX2 (gift of D. Trono; Addgene plasmid #12260) and pseudo typed with pMD2.G (gift of D. Trono; Addgene plasmid #12259). Following transfection, media was changed at 6hrs and 72hrs supernatants were collected and concentrated using ultracentrifugation above a 30% sucrose cushion for 2hrs at 25,000 rpm (Beckman; SW32-Ti). Supernatants were discarded and pelleted lentivirus was re-suspended overnight at 4C in 200 µL of PBS. Sorted PNECs were spin-transduced (800*g for 30’ at 37C) with 10uL of concentrated lentivirus in a total volume of 100uL in the presence of 8ug/mL polybrene (Sigma Aldrich).

#### Intratracheal infections with adenovirus

Mice were anesthetized using inhaled 2.5% isoflurane and infected with 10^6^ pfu Ad5.Spc-Cre or Ad5.Cgrp-Cre (Iowa Viral Vector Core Facility) via an intranasal or intratracheal route. Infections were performed in a total volume of 50 µL where adenoviral concentrates were gently mixed with serum free EMEM and CaCl2 (10mM final) and allowed to salt out at room temperature for 20’ before placing on ice and using within the hour (*87*).

#### Select chemicals and in vivo administration

The following chemicals with their working doses and routes of administration were used for *in vivo* experiments and prepared prior to use: Tamoxifen (200mg/kg IP; Med Chem Express), EdU (100mg/kg IP; Med Chem Express), BrdU (100mg/kg IP; Med Chem Express), naphthalene (275mg/kg IP; Sigma Aldrich), osimertinib free base (25mg/kg PO; LC Labs). Tamoxifen and naphthalene were prepared in corn oil. Naphthalene was administered between the hours of 0800-1000 and mice were placed on Nutra-Gel Complete Nutrition gel cups (Bio-Serv) for 48hrs following administration in addition to standard husbandry. Weight loss peaked 3-4d following naphthalene administrations; however, if mice experienced a decrease >20% their starting body weight following naphthalene our protocol necessated euthanasia and mice were censored from the study. EdU and BrdU were prepared in PBS. Osimertinib was prepared in a vehicle consisting of 30% Kollisolv PEG 300, 1% Tween-80 and 10mM Tris pH 8.0 (Sigma Aldrich). All compounds were re-suspended with gentle sonication in a water bath set to 37C for 1 hour (Branson) and protected from light. Etoposide, palbociclib, MYCi975, and adavosertib were prepared in DMSO (Selleck Chemicals).

#### RNA isolation, cDNA preparation and qPCR

RNAs from 10,000 lineage+ sorted tracheal cells were extracted using a Quick RNA microprep kit (Zymo). 10ng of RNA was then converted to cDNA in a PCR thermocycler using a SuperScript™ VILO™ cDNA synthesis kit. qPCR was performed on an QuantStudio 7 instrument (ABI) using 2X Universal SYBR green fast mix (ABClonal). *Ascl1* expression was normalized to *tdTomato* using the following primer sequences:

*Ascl1* For. 5’-TTCTCCGGTCTCGTCCTACTC-3’ *Ascl1* Rev. 5’-CCAGTTGGTAAAGTCCAGCAG-3’ *tdTom* For. 5’-CCTGTTCCTGGGGCATGG-3’ *tdTom* Rev. 5’-TGATGACGGCCATGTTGTTG-3’

#### Western Blotting

Total proteins were extracted from organoid cell pellets using 1X RIPA buffer (Sigma Aldrich) supplemented with 1X Halt™ protease and phosphatase inhibitor cocktail (Pierce). Lysates were sonicated for 10 pulses at 30% output and 50% duty cycle using a refrigerated Branson 250 Sonifier. Crude extracts were then clarified at 20,000*g for 20’ at 4C using a benchtop refrigerated microcentrifuge (Eppendorf). SDS-PAGE samples were prepared in NuPAGE LDS sample buffer and reducing agent, heated to 90C for 10’, cooled and loaded onto 4-12% Bis-Tris Novex Midi Protein Gels (Thermo Fisher Scientific). Protein gels were transferred onto Immobilon-FL PVDF membranes and blocked using LiCOR TBS Blocking buffer. The following primary antibodies were used at 1:1000 dilution (4C; overnight) in blocking buffer supplemented with 0.05% Tween-20: anti-Myc (CST #9402), anti-pMyc(Thr58) (CST #46650), anti-pChk1(Ser345) (CST #2348), anti-Ncam1 (CST #99746), anti-GFP (CST #2956), anti-pH3(Ser10) (CST #9701), anti-H3 (CST #4499), anti-pH2A.X(Ser139) (CST #9718), anti-pRb1(Ser807/811) (CST #8516), anti-pRb1(Ser780) (CST #8180), anti-Parp (CST #9542), anti-p53 (CST #2524), anti-Ascl1 (BD #24B72D11.1), anti-Dll3 (CST #2483), anti-Bcl2 (CST #3498) and anti-Bim (CST #2933). Secondary conjugated antibodies were used at 1:20,000 (LiCOR) and blocking buffer was supplemented with 0.05% Tween-20 and 0.01% SDS. Membranes were dried prior to imaging on a LiCOR CLx imaging system and images were prepared in ImageStudio (LiCOR).

#### Statistical Tests

Comparative testing between groups was performed in GraphPad Prism software (v9.5.1) and significance thresholds (p-values) per test are indicated throughout figure legends. Paired t-tests assumed Gaussian distribution (*parametric*) and survival analyses were analyzed using a LogRank (Mantel-Cox) test.

Other statistical methodology and comparisons are provided in the ***Supplementary Documentation*** section.

### Single Cell and Bulk Sequencing

#### Single cell RNA-sequencing and bulk ATAC-sequencing sample preparation

All samples were sorted from freshly digested lung tissues. Sample cold time did not exceed 2 hours. Briefly, both male and female mice were collected for these studies, and lungs processed as above. A target of 50,000 tdTomato+, viable cells per animal was sorted and collected in 15cc Falcon tubes containing 5cc of MACS buffer supplemented with 10uM Rock inhibitor Y-27632 (Med Chem Express). Tube were end-over-end mixed for 10’ prior to collecting sorted cells to lubricate the sides of the collecting tubes. Sorted cells were then spun down at 500*g for 3’ in a swinging bucket rotor, re-suspended in the same sorting buffer and manually counted by trypan blue exclusion. Triplicates were pooled for a target total cell number of 60,000 in 50uL to load less than 20uL into the 10X controller or perform TDE1-based tagmentation reactions on small volumes (Illumina Tagment DNA TDE1 Enzyme and Buffer Kit). The 10X Genomics Chromium Platform was used to generate a targeted 5,000 single cell Gel Bead-In-Emulsions (GEMs) per sample, loaded with an average initial cell viability of >80%. scRNA-seq libraries were prepared following the 10X Genomics user guide (Chromium Next GEM Single Cell 3’ Reagent Kits, v3.1 Dual Index Chemistry; 10X Genomics). ATAC-seq libraries were prepared using established protocols (*88*). After encapsulation, emulsions were transferred to a thermal cycler for reverse transcription (RT) at 53°C for 45 min, followed by heat inactivation for 5 min at 85°C. cDNA from the RT reaction was purified using DynaBeads MyOne Silane Beads (Thermo Fisher Scientific) and amplified for 12 cycles using Amplification mix and primers provided in the Single Cell 3’ reagents module 1 (10X Genomics).

After purification with 0.6X SPRI beads (Beckman Coulter), cDNA quality and yield were evaluated using Agilent Bioanalyzer with a 25-1000bp DNA standard. Using a fragmentation enzyme blend (10X Genomics) the libraries were fragmented, end-repaired and A-tailed. Products were double side cleaned using 0.6X and 0.8X SPRI beads, and adaptors provided in the kit were ligated for 15 min at 30°C. After cleaning ligation products, libraries were amplified and indexed with unique sample index i7 through PCR amplification. The number of PCR cycles was chosen based on cDNA yield for each sample individually and did not exceed recommended thresholds. Final libraries were cleaned using 0.6X then 0.8X SPRIselect beads and their quality and size were evaluated using a Bioanalyzer. Bulk ATAC samples were processed like the above, with a target pooled cell number of ∼50,000. All single cell RNA and bulk ATAC library preparations and library QC were carried out by the WCM Epigenomics core facility. Libraries were pooled and sequenced on a HiSeq2500 or NovaSeq instrument (Illumina) at the WCM NGS core facility. Flow cell paired-end reads per samples followed recommendations in the 10X Genomics guide or established protocols for Omni-ATAC (*88*). Illumina’s demultiplexing pipeline to generate a tar archive of gzipped FASTQ files for each sample.

### Computational Analyses

Single cell RNA-seq data were mapped with CellRanger (v5.0.1) and processed with custom Python scripts for data pre-processing, cell type annotations, lineage probability assignments and analyses, as well as cell state trajectory inference, which are fully described in supplementary **notes S1** to **S4**. Bulk ATAC-seq analyses, including quality control and pre-processing, are fully described in supplementary **note S5**.

### Single cell RNA-seq Data Visualization

#### Two-dimensional embeddings

The global atlas of all cells in the sequenced in the study, including tumor, stroma, lymphoid and myeloid subsets were visualized using a Uniform Manifold Approximation and Projection (UMAP) (*see **Supplementary Documentation***), given the diversity of cell types represented. The global UMAP was computed using *k=20* nearest neighbors, *min_dist=0.25*, and remaining default *scanpy* (*89*) parameters. Force-directed graph layouts (FDLs) were used to visualize the tumor-epithelial cell subset because these better capture cell state transitions and more continuous cell state relationships on the cell-cell similarity graph. The FDLs were computed using *k=25* nearest neighbors for the original 24-sample dataset (**Fig. 2** and **fig. S1, S4,** and **S11**) and *k=30* nearest neighbors for the extended 27-sample merged dataset (**fig. S15A**), with remaining default *scanpy* parameters (*see **Supplementary Documentation***).

#### Gene expression along the LUAD to SCLC trajectory

The transitional highly variable gene (HVG) heatmap was generated using the *CellRank* heatmap plotting function, which uses a generalized additive model (GAM) to smooth expression along the given trajectory. Imputed expression was used to generate this visual, and expression was normalized to a range of [0,1]. The order of genes was determined by the expression peak along the terminal state probability continuum. Transition genes are defined as HVGs without an expression peak at either end point of the inferred terminal state probability. Kernel density estimates (KDEs) above the heatmap were generated by ranking the given variable along the given trajectory. Similarly, gene trend curves were generated using the built-in plotting method provided by *CellRank* using the same GAM method and min-max normalization as in the heatmap visual described above.

#### Additional Information

Additional information regarding methods can be found in the **supplementary notes**. For scRNA-seq data pre-processing, see **note S1**; for macro cell type annotations and tumor-epithelial subset selection, see **note S2**; for differential gene expression and pathway enrichment, see **note S3**; for single cell trajectory analysis, see **note S4**; for bulk ATAC-seq analyses, see **note S5**.

---

**Supplemental Figure 1.**
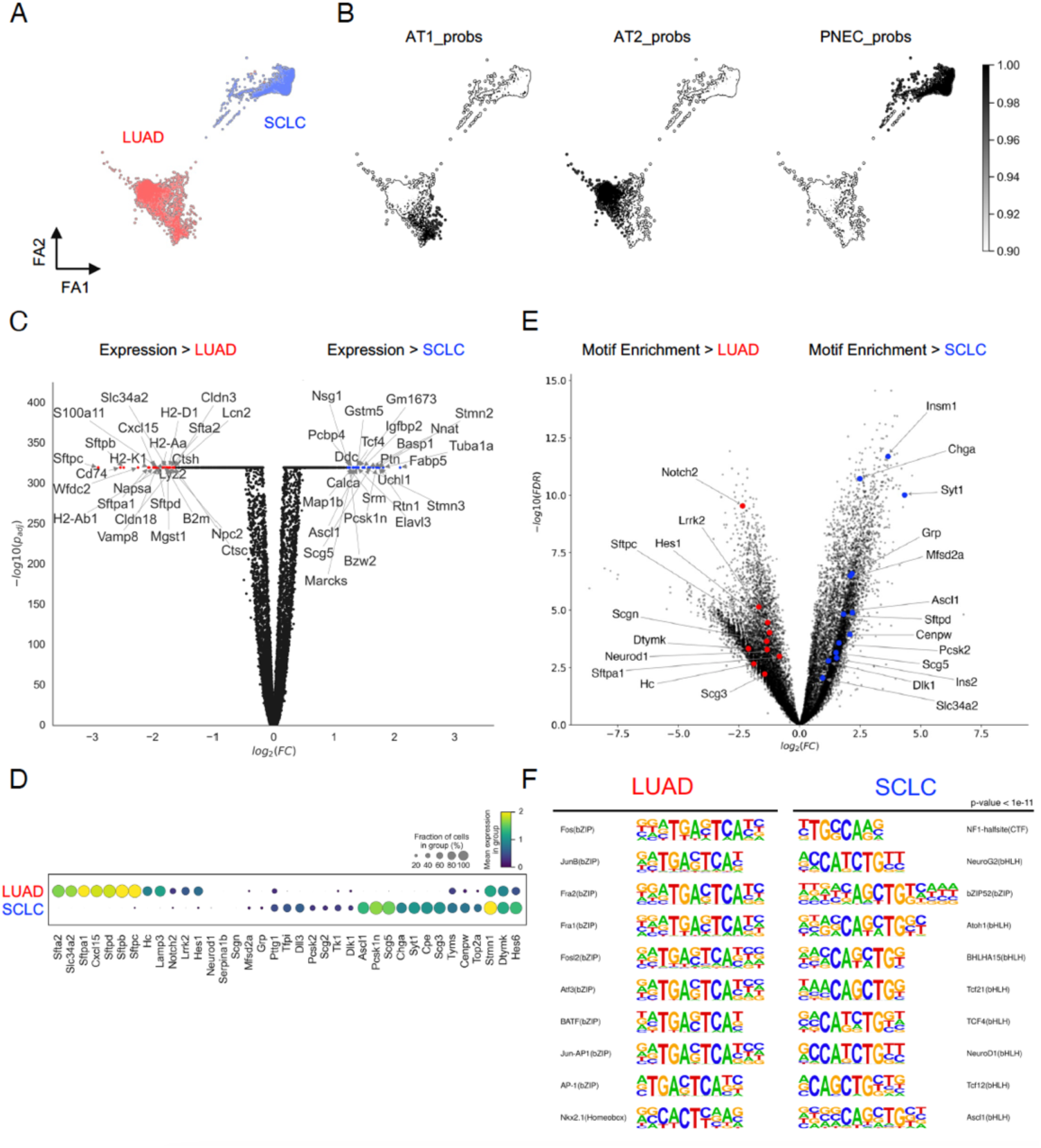
Characterization of LUAD and SCLC transcriptional programs in the ERPMT model. A) Force-directed layout (FDL) (*91*) of all tumor-epithelial cells (tdTom+) derived from the LUAD and SCLC models (as in **Fig. 1D**), colored by source (*A*) and lineage probability (*see Methods*) (*B*) (*92*). C) Volcano plot showing differentially expression genes (DEGs, *see Methods*) in the ERPMT-derived SCLC vs. LUAD models; top 25 DEGs per model are labeled (**table S2**). D) Dot plot showing frequency of expressing cells (*node size*) and log-transformed average expression (*node color*) of AT2 and PNEC lineage genes (**Fig. 1D**) stratified by model. E) Volcano plot showing differentially accessible peaks for SCLC vs. LUAD models (n = 3 LUAD and n = 2 SCLC biological replicates; **table S2**). Peaks are annotated by the nearest gene and aggregated by significance. Significant peaks (FDR < 0.01) associated with AT2 and PNEC lineage markers (or *Insm1*) are labeled, and scatter points are colored by model. F) Enriched motifs associated with differentially accessible (DA) peaks (FDR < 0.01, abs(log2FC) > 1) between LUAD and SCLC models. The top 10 most significant known motifs present in at least 10% of the DA peaks for each model are shown.

**Supplemental Figure 2.**
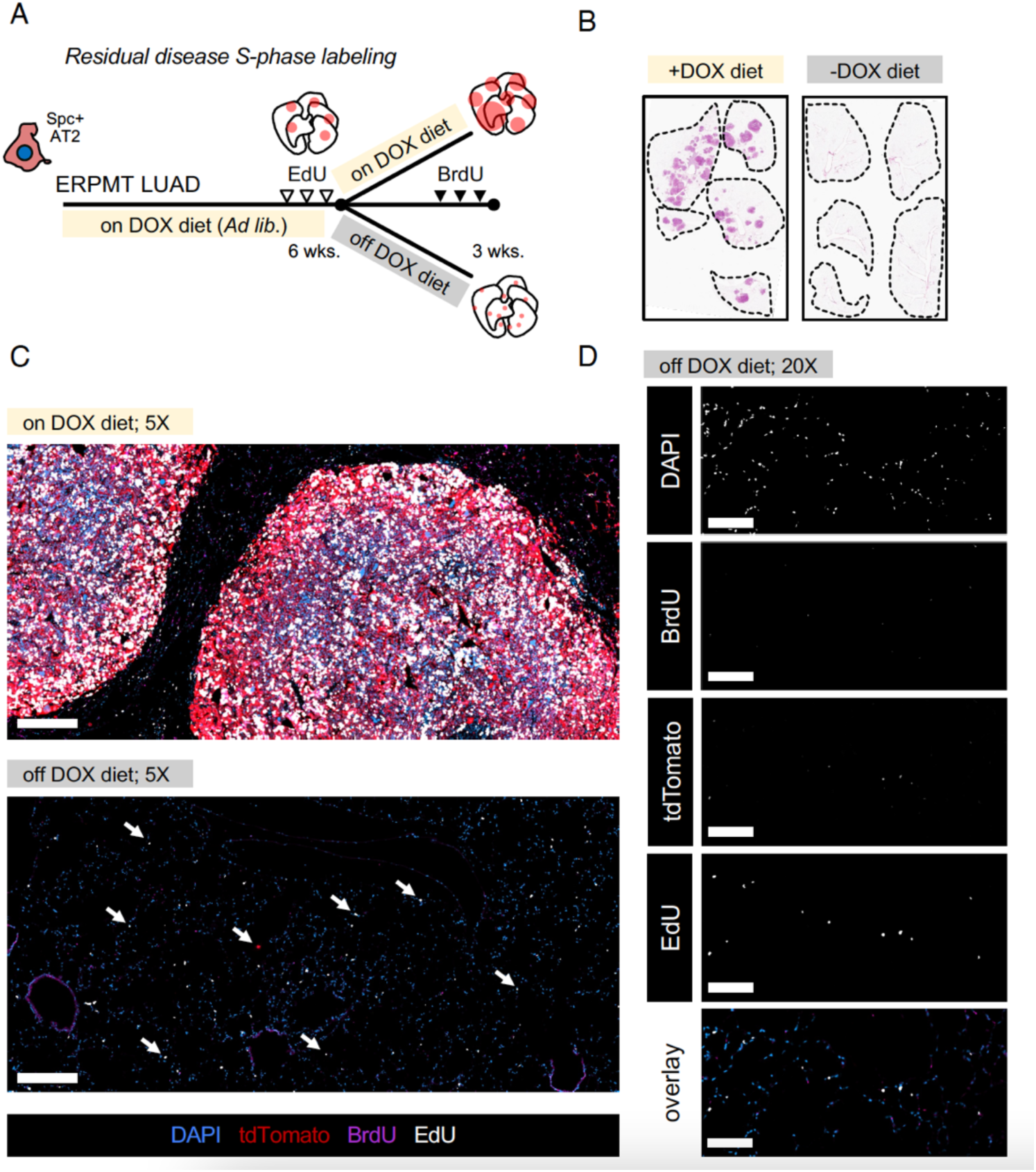
Following EGFR withdrawal, residual tumor cells are not highly proliferative. A) Sequential EdU / BrdU labeling strategy in ERPMT LUAD mice on study to label and track cycling cells. B) Representative sagittal H&E sections from mice in each group at the end of study (week 9; n = 5 per group). Immunofluorescence (IF) section of tumor lesions in mice from each experiment group. Arrows in bottom IF panel highlight residual, single cell spacing across a large airway section and +/- label uptake; 500um scalebar. C) Higher magnification of lung section from mice off DOX diet group demonstrating high frequency of tdTom+ / EdU+ / BrdU- cells; 100uM scalebar (*overlay as in C*).

**Supplemental Figure 3.**
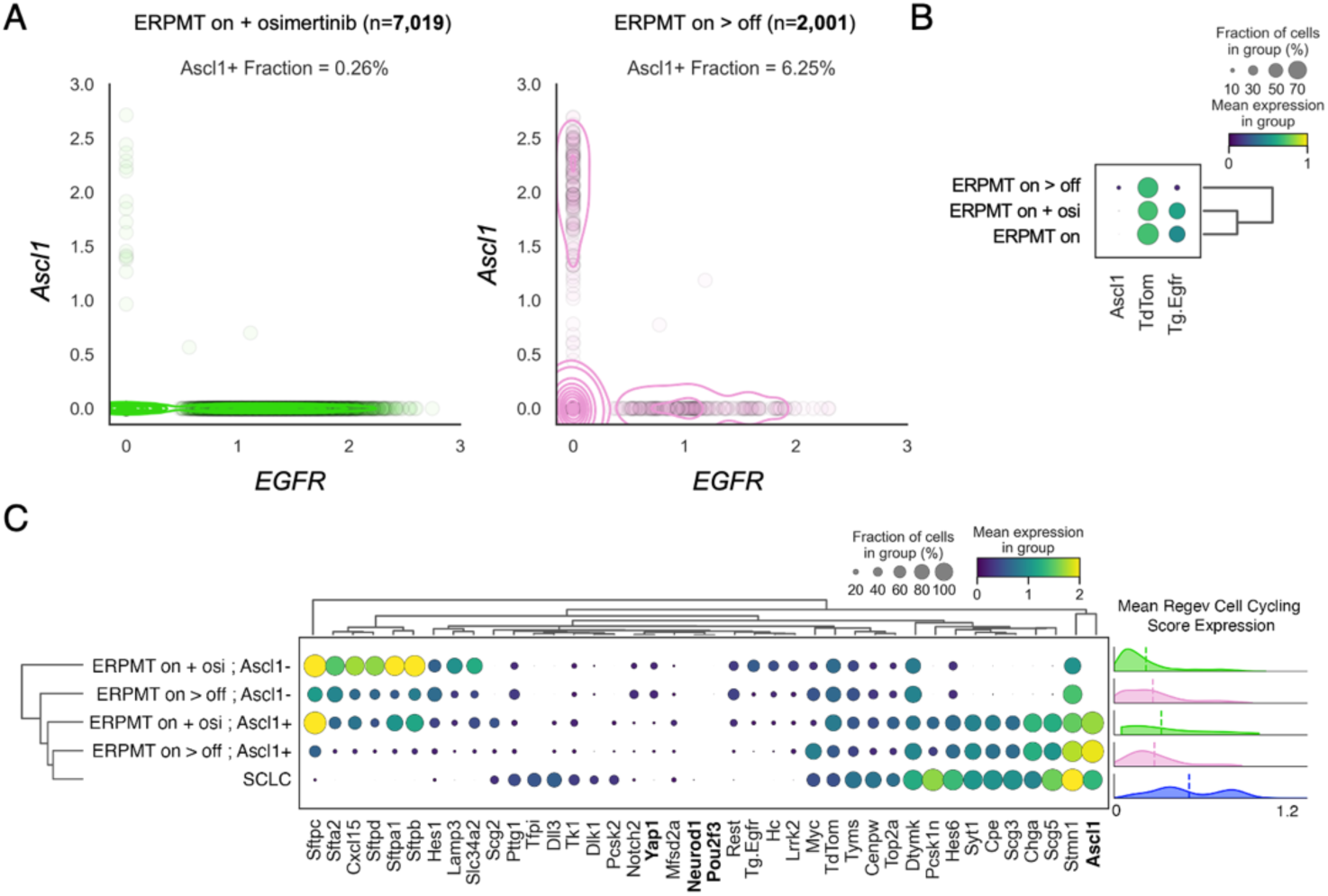
E*G*FR extinction is required for *Ascl1* expression. A) Scatter plot of log-transformed expression of *EGFR* and *Ascl1* in single tdTom+ cells isolated from 1mo residuals in the ERPMT LUAD model after *EGFR* was withdrawn pharmacologically (mice treated with osimertinib on DOX; *green*) or genetically (off DOX; *pink*). Frequency of *Ascl1*-positive cells (log-transformed expression > 0) and total number of cells sequenced listed above. B) Dot plot showing frequency of expressing cells (*node size*) and log-transformed expression (*node color*) of *EGFR, Ascl1 and tdTom* across select LUAD single cell datasets. C) Log-transformed average expression of AT2 and PNEC lineage markers, model oncogenic drivers (*Myc* and *Tg.EGFR)* and SCLC markers (*NeuroD1, Pou2f3,* and *Yap1*) (*8*) expressed in at least 5% of cells, grouped by sample. The ERPMT off DOX and ERPMT on DOX + osimertinib samples were further stratified by *Ascl1* expression status. Genes and sample groups are clustered using the centroid Euclidean distance method. *Right*, kernel density estimates showing mean imputed expression of a cell cycling signature (*38*) across samples; dashed line denotes the distribution median.

**Supplemental Figure 4.**
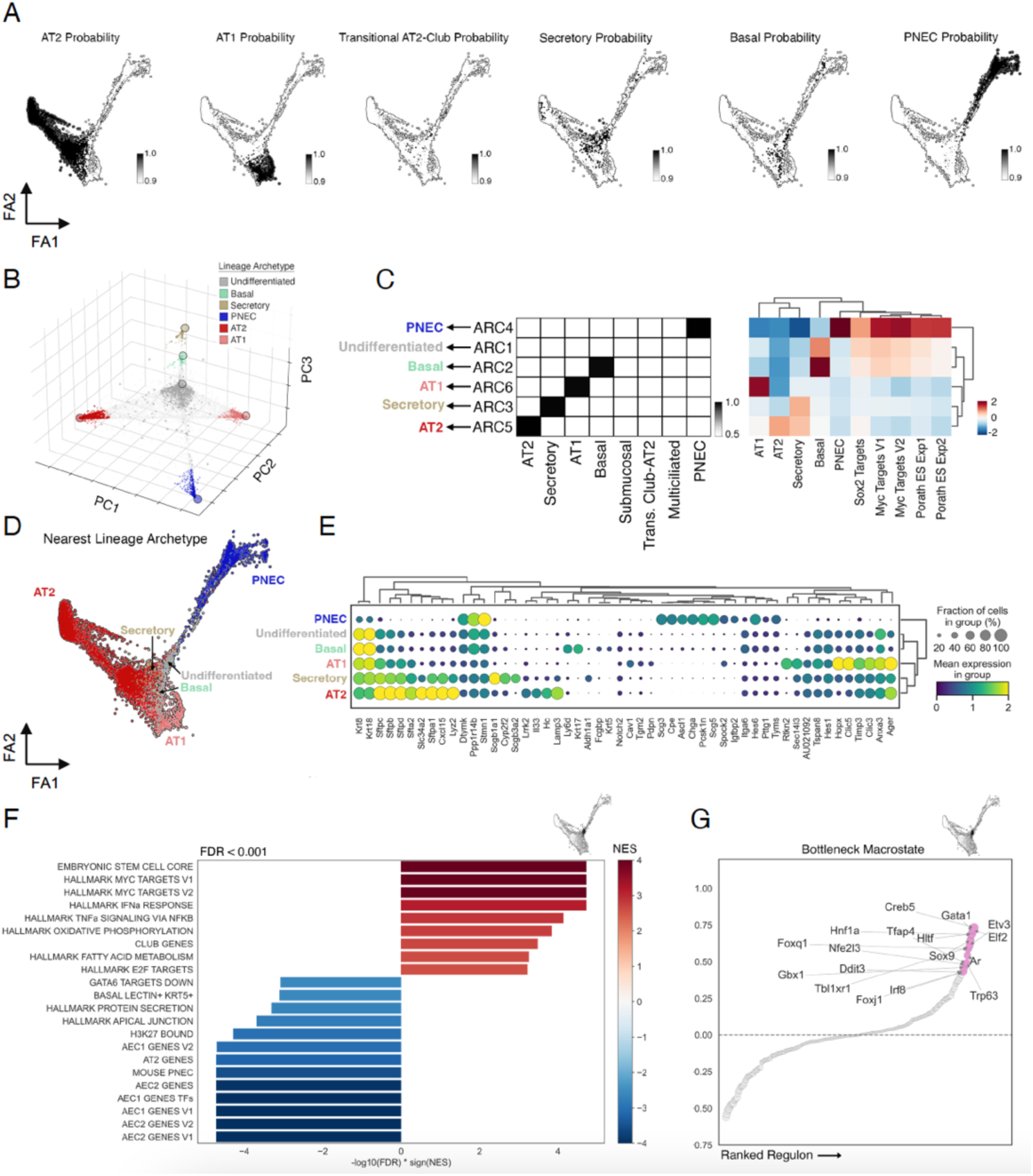
Characterizing alveolar-derived LUAD bottleneck on the transition to SCLC. A) Force-directed layout of cell states captured along the transition from at AT2 cell to ERPMT LUAD and finally towards a neuroendocrine fate (as in **Fig. 2E**) colored by lineage probability. B) PCA projection of the lung epithelial lineage probability per cell colored by lineage archetype soft cluster (*see Methods*); archetypes overlaid. C) Heatmap showing average lineage probability (*left*) and mean imputed expression (*z-normalized*) of marker genes for each lung epithelial lineage (**table S1**), as well as Myc and stem cell-related gene signatures, for all cells in each archetype soft cluster (*right*) (**table S3**). Rows are clustered based on the Manhattan distance, and columns are ordered manually (*left*) or clustered based on the Manhattan distance (*right*). Assignment of archetypes to lung epithelial lineages shown with arrows. D) FDL in (A) colored by nearest lineage archetype. E) Dot plot showing frequency of expressing cells (*node size*) and log-transformed mean expression (*node color*) of lineage marker genes differentially expressed (Bonferroni p-value < 0.05, abs(log2FC) > 0.05, fraction expressing >20%) in at least one archetype soft cluster. Genes are cluster by Pearson correlation distance and archetypes are clustered by Euclidean distance. F) Bar plots showing top pathways (FDR < 0.001) differentially expressed along bottleneck macrostate probability (FDL; *top right*). The x-axis shows -log10(FDR) times the sign of the normalized enrichment score (NES) and bars are colored by NES. G) *SCENIC* (*93, 94*) transcriptional regulatory modules (“*regulons*”, *see Methods*) correlated with the bottleneck macrostate probability. The top 5% (n = 17) of correlated regulons are labeled (**table S1**).

**Supplemental Figure 5.**
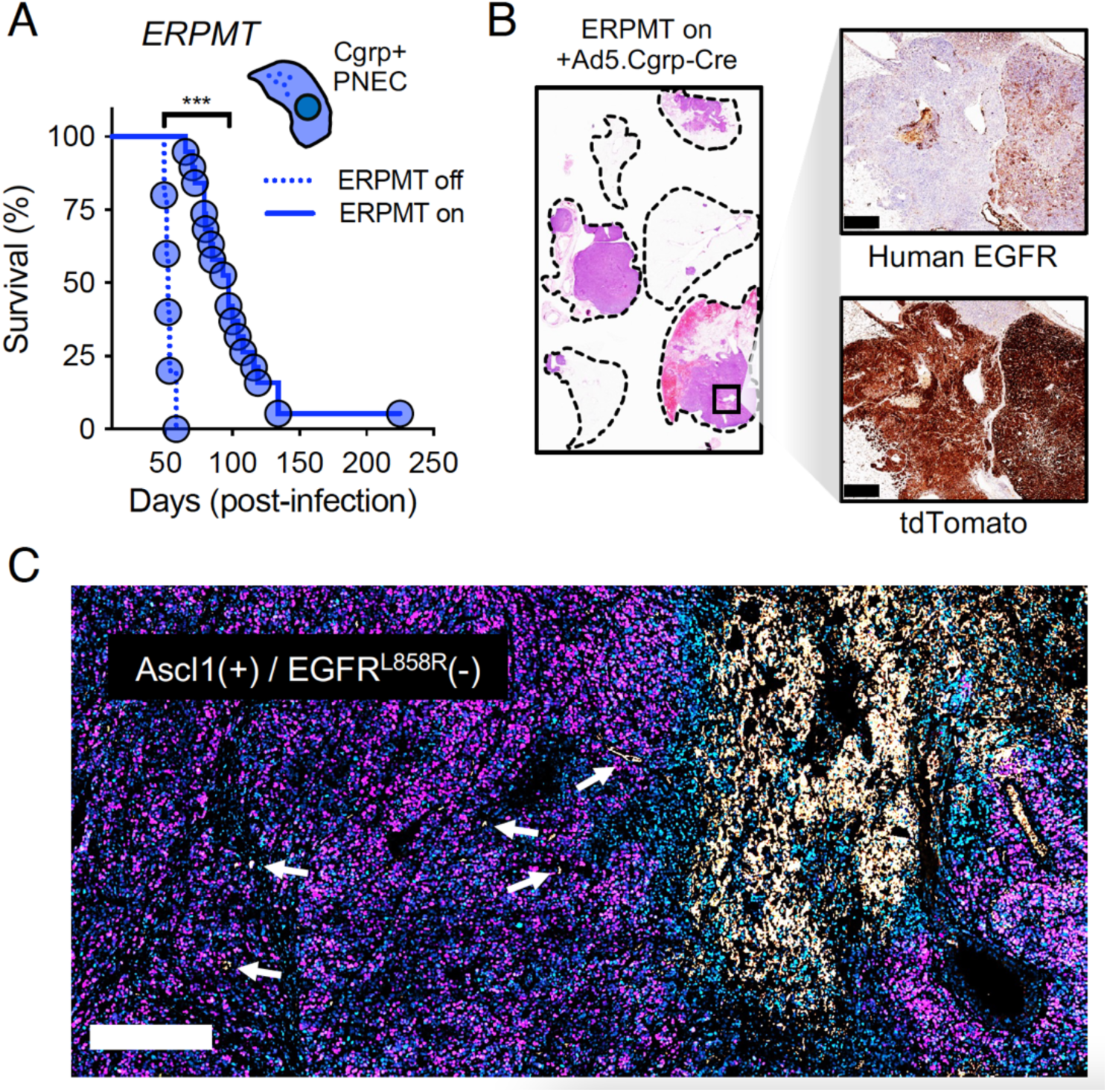
PNEC cells are refractory to generating LUAD in the ERPMT model. A) Initiating the ERPMT model in a neuroendocrine cell of origin (Ad5.Cgrp-Cre) and survival of mice on or off DOX. Dashed blue line is off DOX (n = 5) and solid blue line is on DOX diet from start of study (n = 19); ***p<0.001. B) Representative histology from mice in the “ERPMT on” group in *A* with zoomed in boxed tumor section showing human EGFR and tdTomato IHC staining; 1mm scalebar. C) Immunofluorescence for EGFR^L858R^ (*yellow*) and Ascl1 (*purple*) demonstrating staining exclusivity in tumor section from *B*; 100um scalebar, DAPI+ nuclei in *blue*. Select EGFR+ cells within the larger Ascl1-positive / EGFR-negative field are indicated with white arrows.

**Supplemental Figure 6.**
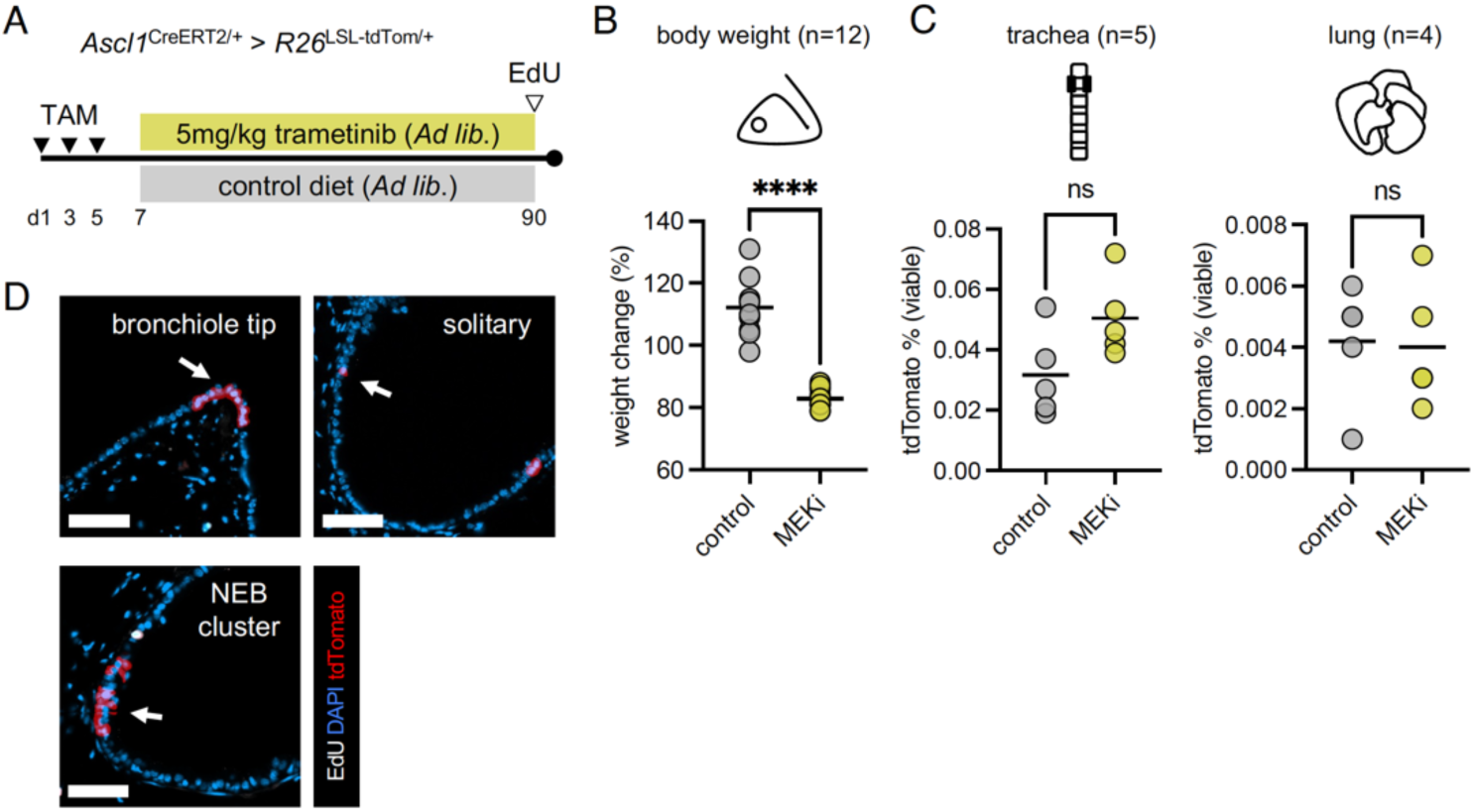
MEK chemical inhibition does not increase PNEC frequency, anatomic location, or proliferative status. A) Labeling timeline to determine the effects of systemic MEK inhibition (MEKi) on Ascl1-lineage traced cells (trametinib formulated at 5mg/kg in chow provided *ad libitum*). B) Percent change in animal body weight from point of randomization of study. Protocol limits necessitated termination of study at ∼3mo on trametinib diet due to weight loss and malaise (n = 12 per arm); ****p<0.0001.C) Quantitation of tdTomato+ cells from single cell enzymatic digestions of the tracheas or lungs from mice on study for 3mo; n = 4-5 per arm. D) Representative immunofluorescence images of distinct anatomic locations of lineage labeled PNECs throughout the airway in mice at 3mo post-labeling on MEKi diet; NEB = neuroepithelial body (50um scalebar).

**Supplemental Figure 7.**
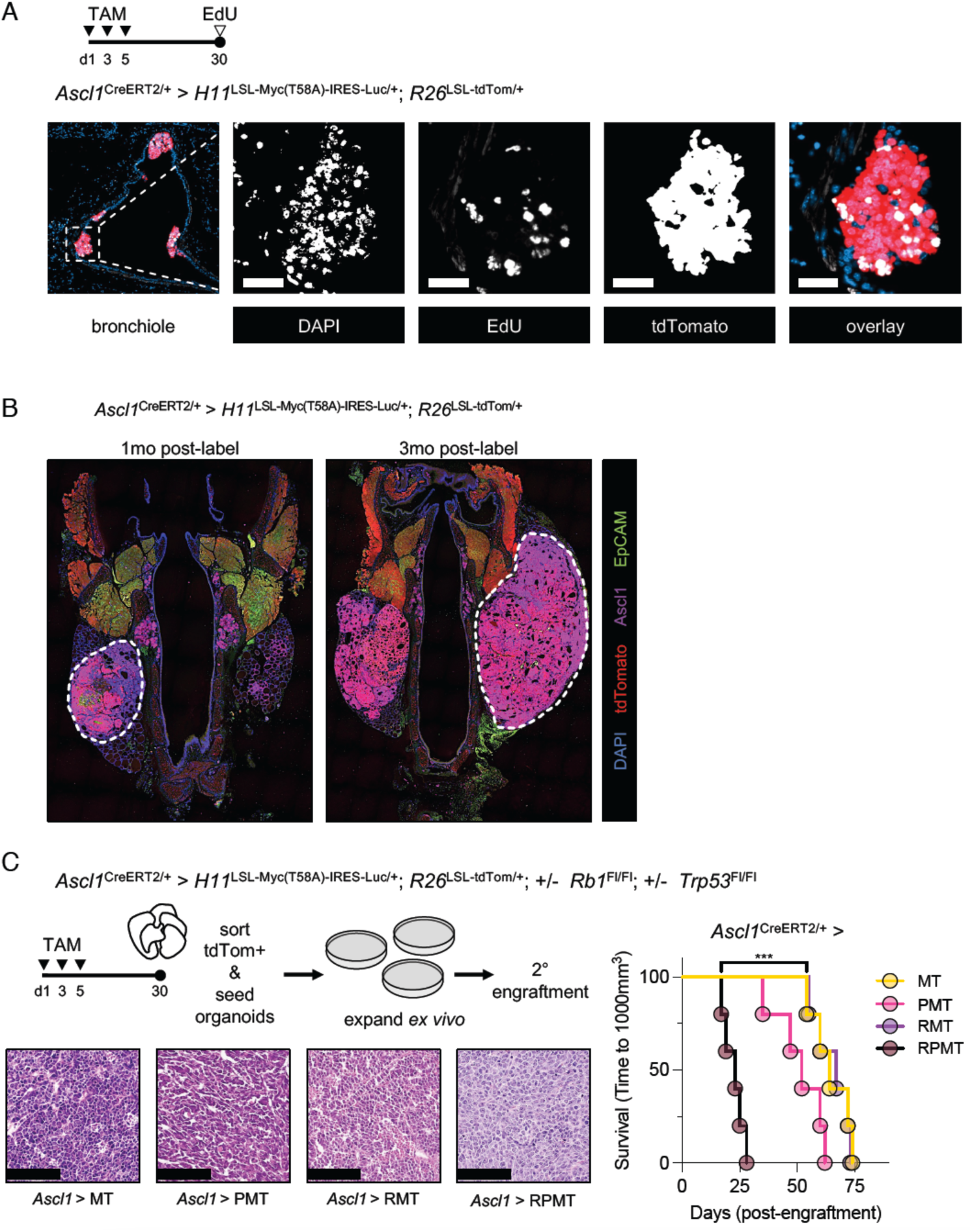
Myc is sufficient to transform the *Ascl1*+ lineage. A) Experimental outline and representative immunofluorescence of bronchioles in *Ascl1*>MT mice 1mo following tamoxifen labeling. Zoomed in section on one of several proliferative PNEC clusters; 50um scalebar. A single dose of EdU was administered 2hrs before collecting tissues. B) Indirect immunofluorescence staining for indicated targets on tracheal sections from *Ascl1*>MT mice at 1mo and 3mo post-labeling with tamoxifen. Tumor lesions arising within the Ascl1+ thyroid C-cells are outlined in dashed white lines. Sagittal tracheal sections are at the same scale for comparative purposes. C) Sorting and expanding PNECs from the lungs of mice with various genotypes, followed by subcutaneous engraftment into immunocompromised mice. H&E of representative tumors from mice at study endpoint; 200um scalebar. Survival of mice engrafted with lung organoids (athymic/nude; n = 5 per arm). *P* = *Trp53*^Fl/Fl^; *R* = *Rb1*^Fl/Fl^ . Events were recorded as time to reach ∼1000mm^3^ following engraftment of 10^6^ viable cells from organoid cultures; ***p<0.001.

**Supplemental Figure 8.**
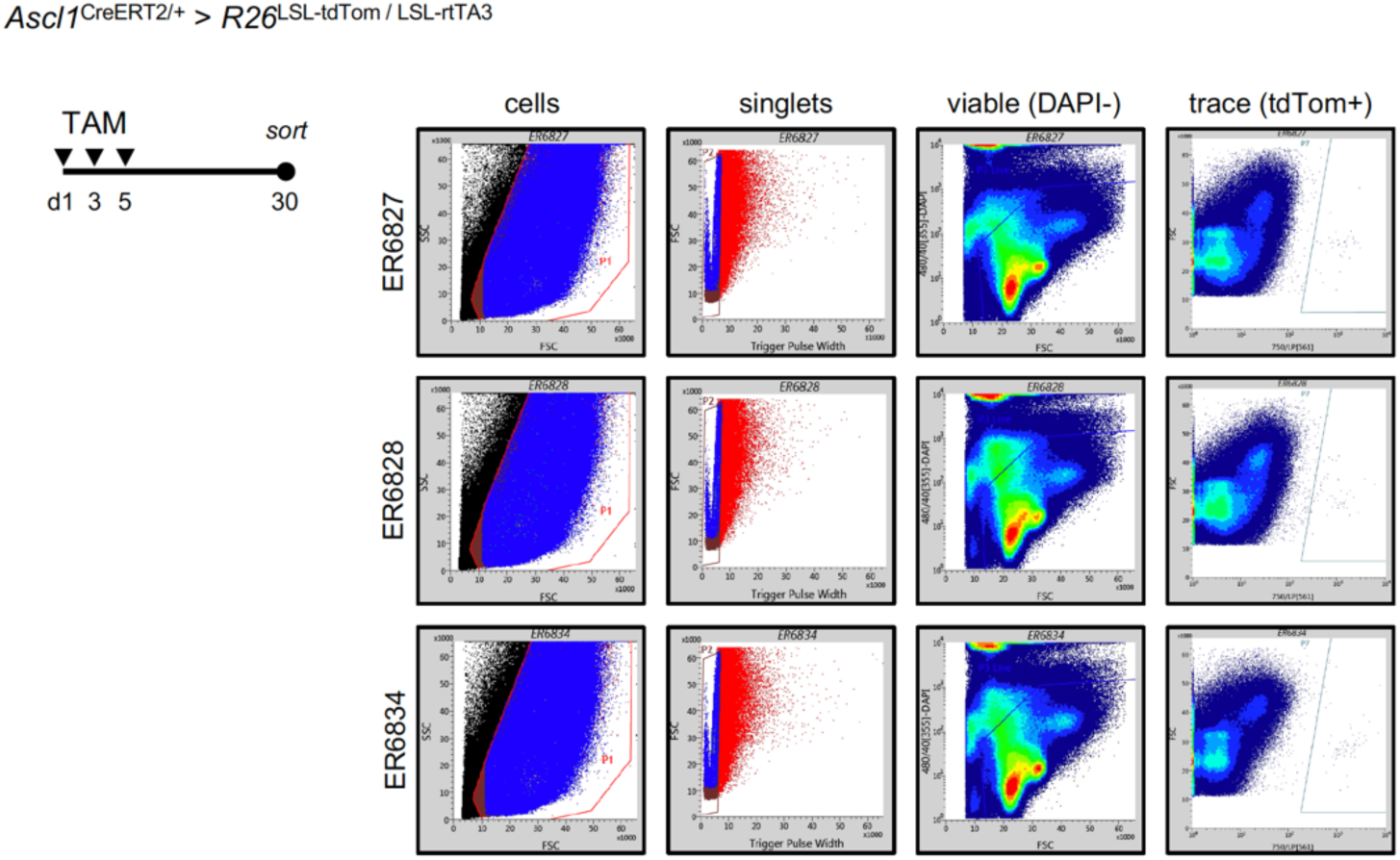
Gating strategy for isolation of pulmonary *Ascl1*^CreERT2^; *R26*^LSL-tdTom/LSL-rtTA3^ cells from single cell digestions. Gating strategy on BD Influx for three separate *Ascl1*^CreERT2^>*Rosa26*^LSL-tdTom/LSL-rtTA3^ male, adult mice at 1mo post-TAM; labeling timeline shown at *left*. The intensity of the Ai14 *Rosa26*^LSL-tdTom^ expression necessitated gating off the far-red channel. Per 2*10^6^ events, the frequencies were as follows: ER6827 (n = 42 cells), ER6828 (n = 59 cells) and ER6834 (n = 65 cells).

**Supplemental Figure 9.**
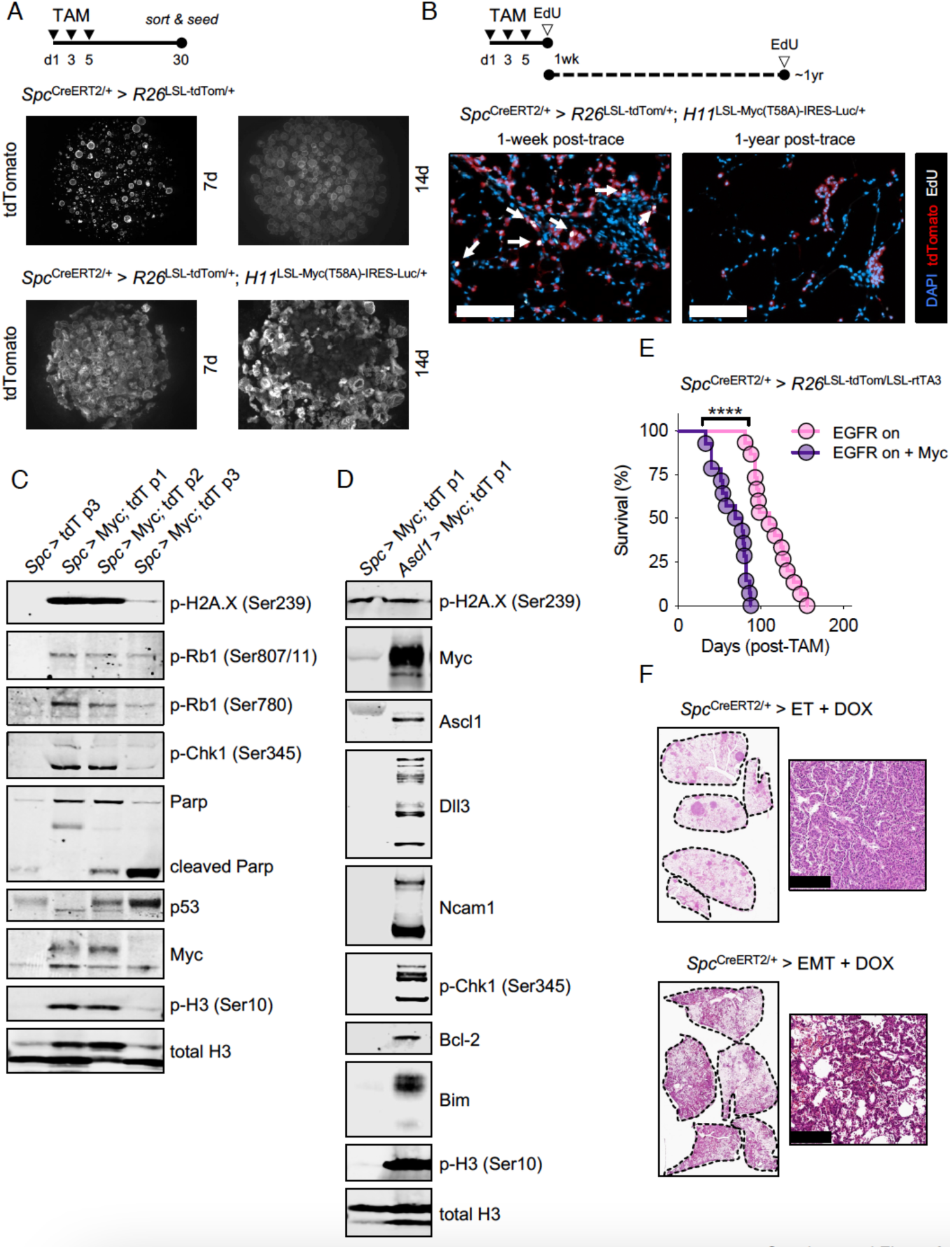
Myc is lethal to AT2 cells, causes replication stress and is relieved by oncogenic EGFR. A) Appearance of spheroid cultures from normal AT2 (Spc>T; *left*) or AT2 cells expressing Myc (Spc>MT; *right*) sorted from mice 1mo post-labeling at various times in culture; *inset*: appearance of Spc>MT spheroid cultures breaking through matrix at 14d after seeding. Organoid cultures were established by seeding 10K sorted cells in 50uL droplets of 50% Matrigel®. Media was changed twice per week. B) Representative IF sections of lineage traced AT2 cells + Myc at one-week and one-year following labeling; 50um scalebar. Labeling schedule provided above; EdU administered once, 2hrs prior to tissue collection. Arrows at *left* designate EdU+, tdTom+ cells. C) Western Blot for markers of DNA damage, replication stress and apoptosis in AT2 organoid cultures +/- Myc and loss of Myc expression over sequential passages. D) Western blot for select neuroendocrine markers, apoptotic proteins and Myc in the first passage of Spc>Myc;tdTom or Ascl1>Myc;tdTom organoids. The band observed in the Ascl1 blot from the Spc>Myc;tdTom sample is suspected to be non-specific. E) Survival of mice where oncogenic EGFR^L858R^ was expressed from a *Spc*^CreERT2^ lineage trace allele alone (n = 15) or in combination with Myc (n = 14); all mice were on DOX diet throughout study. F) Histologic appearance of whole lung sections and select regions at high magnification for mice collected from study in *D*; 200um scalebar.

**Supplemental Figure 10.**
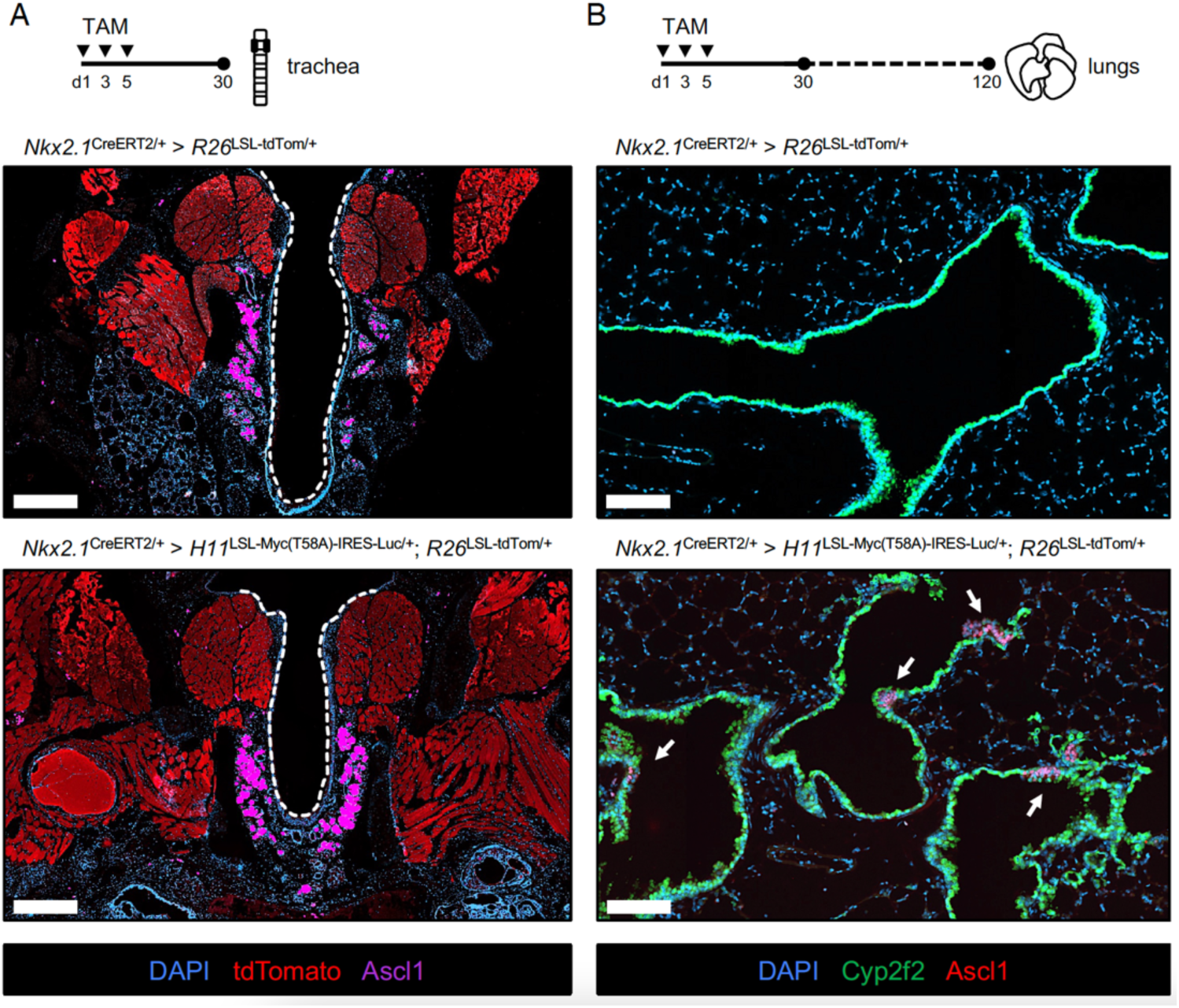
Generalized, conditional Myc expression throughout the airway expands neuroendocrine cells in long-term traces. A) Representative immunofluorescence of sagittal tracheal sections of *Nkx2.1*^CreERT2^-traced cells expressing tdTom (*Nkx2.1*>T*; left*) or with Myc (*Nkx2.1*>MT; *right*) at 1mo post-labeling; 500um scalebar. B) Representative immunofluorescence of bronchi and surrounding lung cells in *Nkx2.1*>T or *Nkx2.1*>MT mice at 4mo post-trace; 100um scalebar. Ascl1+ (*purple*) clusters of cells are indicated using white arrows. Cyp2f2 (*green*) is being used as a secretory cell lineage marker and was consistent with Scgb1a1 staining. Ascl1+ clusters of cells were not observed in time matched *Nkx2.1*>T lungs.

**Supplemental Figure 11.**
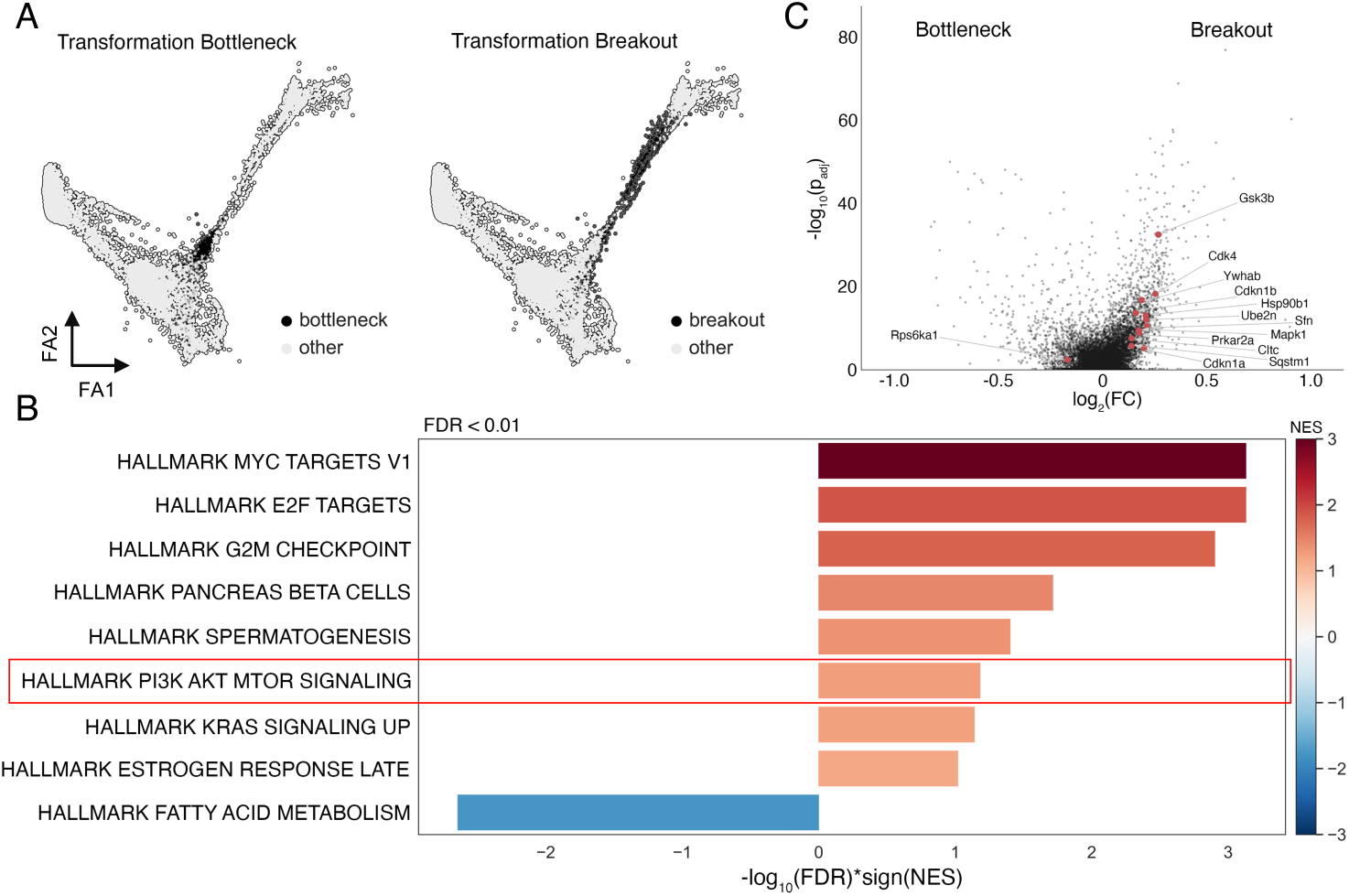
Characterization of tumors cells at the bottleneck and breaking out towards neuroendocrine transformation. A) Force-directed layout of subgroups of tumor cells characterizing the HT dataset bottleneck or breaking through towards a neuroendocrine state (“breakout”). B) Hallmark gene sets differentially enriched (FDR < 0.01) in the bottleneck (left) and breakout (*right*) subpopulations. C) Differentially expressed genes in the bottleneck vs breakout populations, top ranked genes (abs(logFC) > 0.1, Bonferroni p-adjusted < 1e-5) from the Hallmark PI(3)K AKT MTOR signaling pathway are labeled.

**Supplemental Figure 12.**
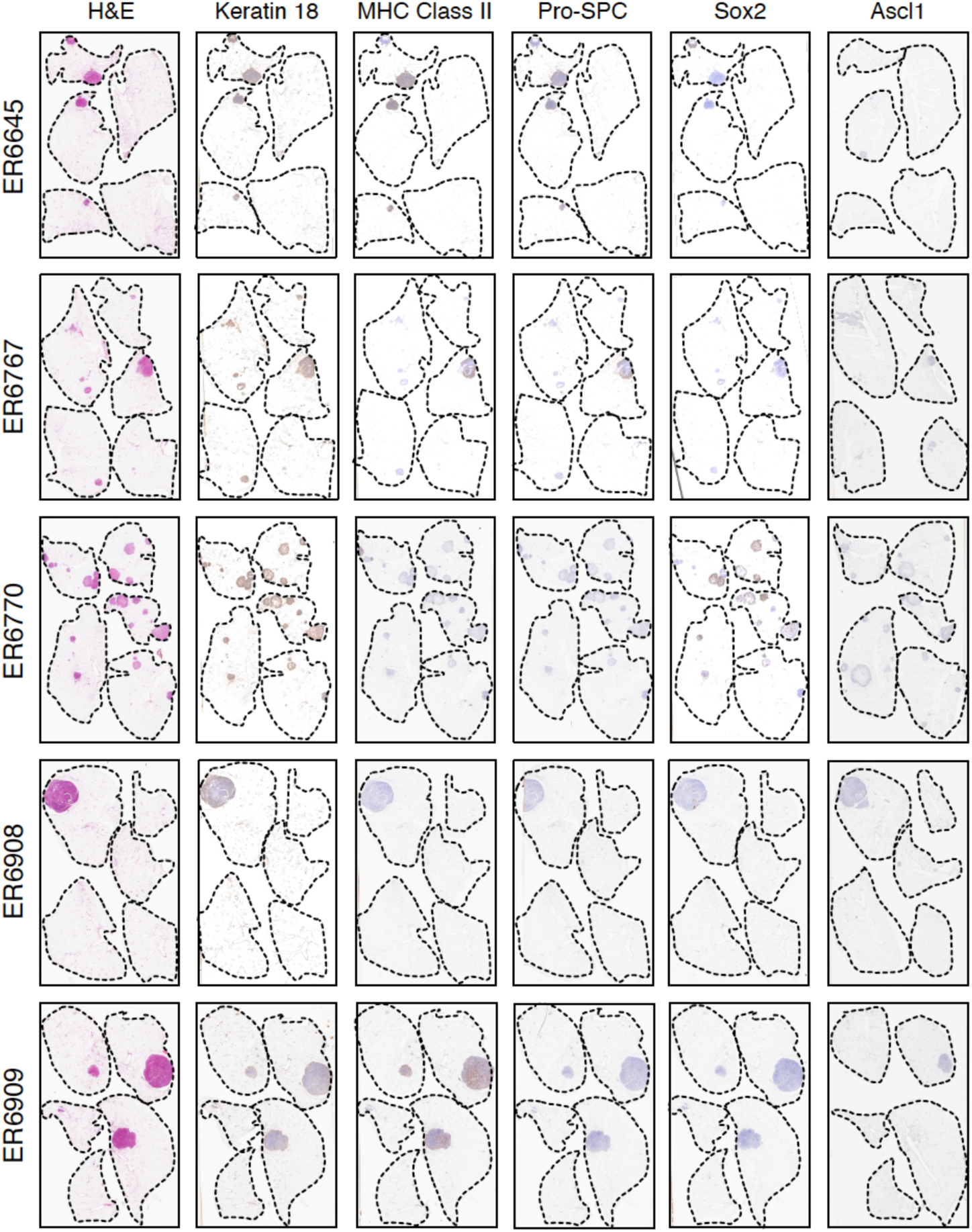
Characterization of Myc-driven tumors that arise from the AT2 lineage. H&E and IHC-based staining for indicated markers across 5 separate *Spc*>Pten;Myc animals (ERXXXX nomenclature) collected at 3mo on study following the sequential administration of tamoxifen as described before. Ascl1 IHC staining was performed on sections cut after ∼50um of depth was cut for unstained slides, and thus the lung margins (*black dotted outlines*) have shifted substantially versus the other slides.

**Supplemental Figure 13.**
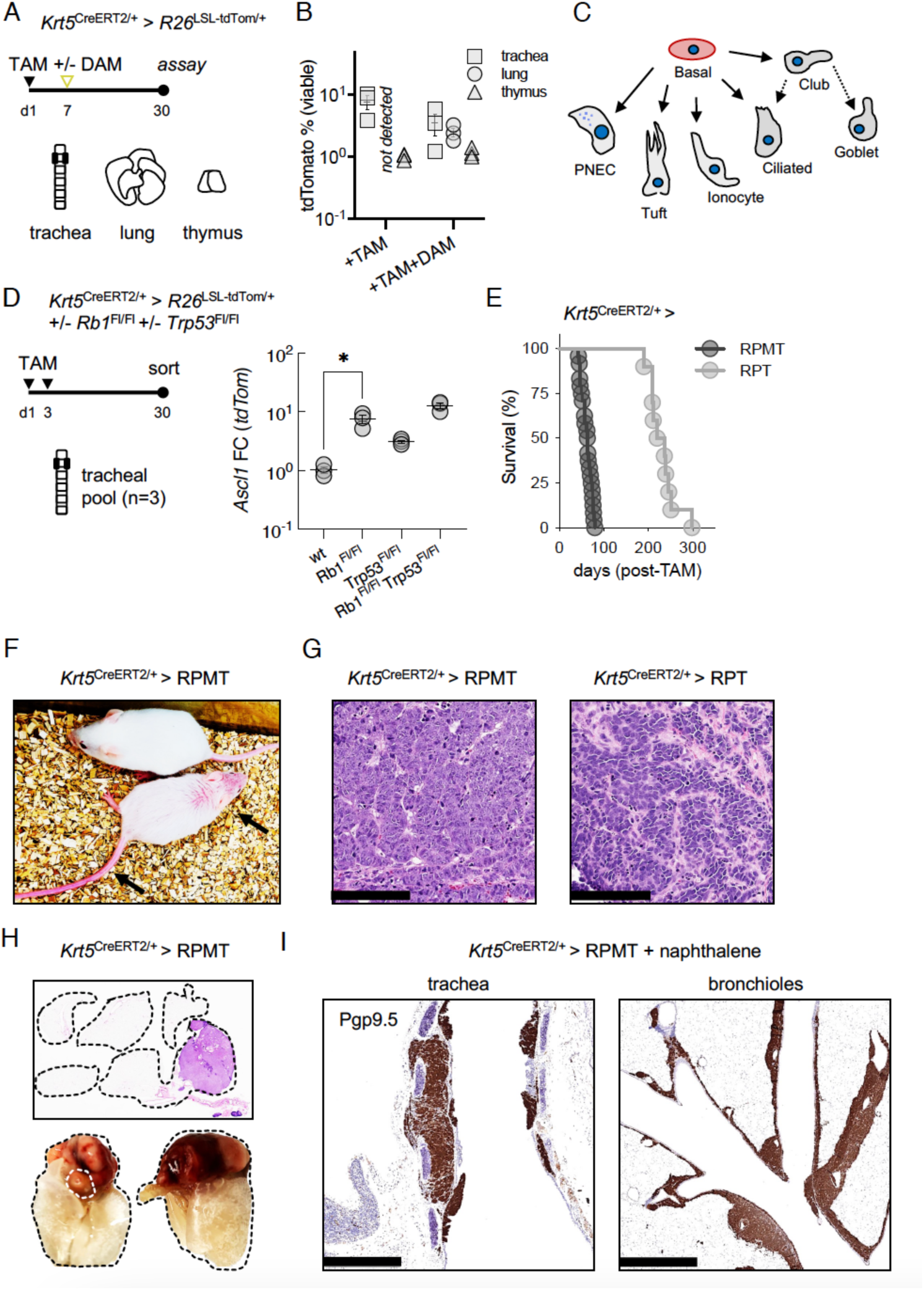
Basal-to-neuroendocrine differentiation upon *Rb1* loss. A) Tamoxifen labeling and naphthalene damage strategy for detection of tdTom+ cells from select tissues in the *Krt5*^CreERT2^ lineage trace model. Mice were administered a single dose of tamoxifen (200mg/kg; IP) on d1 of study and then +/- a single dose of naphthalene (275mg/kg IP; abbreviated as “DAM”) or corn oil between 8AM – 10AM on day 7 of study (n = 3 per arm; +/- SEM). B) TdTom% was not detected in the lungs of labeled basal trace mice without naphthalene damage. Labeling and sorting timeline to isolate tdTom+ cells from the trachea with various genotypes; 3 tracheas were pooled per genotype. C) Cartoon depicting differentiation trajectories (*53*) for pulmonary basal cells. D) *Ascl1* expression fold-change (FC) normalized to *tdTom* by qPCR. E) Survival of mice following *Krt5*-initiated trace of RPT (n = 10) or RPMT genotypes (n = 24). F) Appearance of *Krt5*>RPMT mouse one month following tamoxifen exposure (∼200uL of 20mg/mL IP * 3 doses); arrows calling out expression of tdTom, as compared to an unlabeled control animal (*above*). G) Histologic appearance of tumors arising in *Krt5*>RPT or *Krt5*>RPMT mice when moribund; 100um scalebar. H) Evidence that tumor lesions involve the thymus in *Krt5*^CreERT2^ initiated SCLC models. Whole lung H&E with example of cleared lung tissues below outlined in black dashed line; white dashed line outlines heart, rotating fixed tissue 180°. I) IHC staining for Pgp9.5 in *Krt5*>RPMT tracheal (500um scalebar) and bronchiole (1mm scalebar) sections two months following naphthalene damage where tamoxifen was administered one week prior to damage.

**Supplemental Figure 14.**
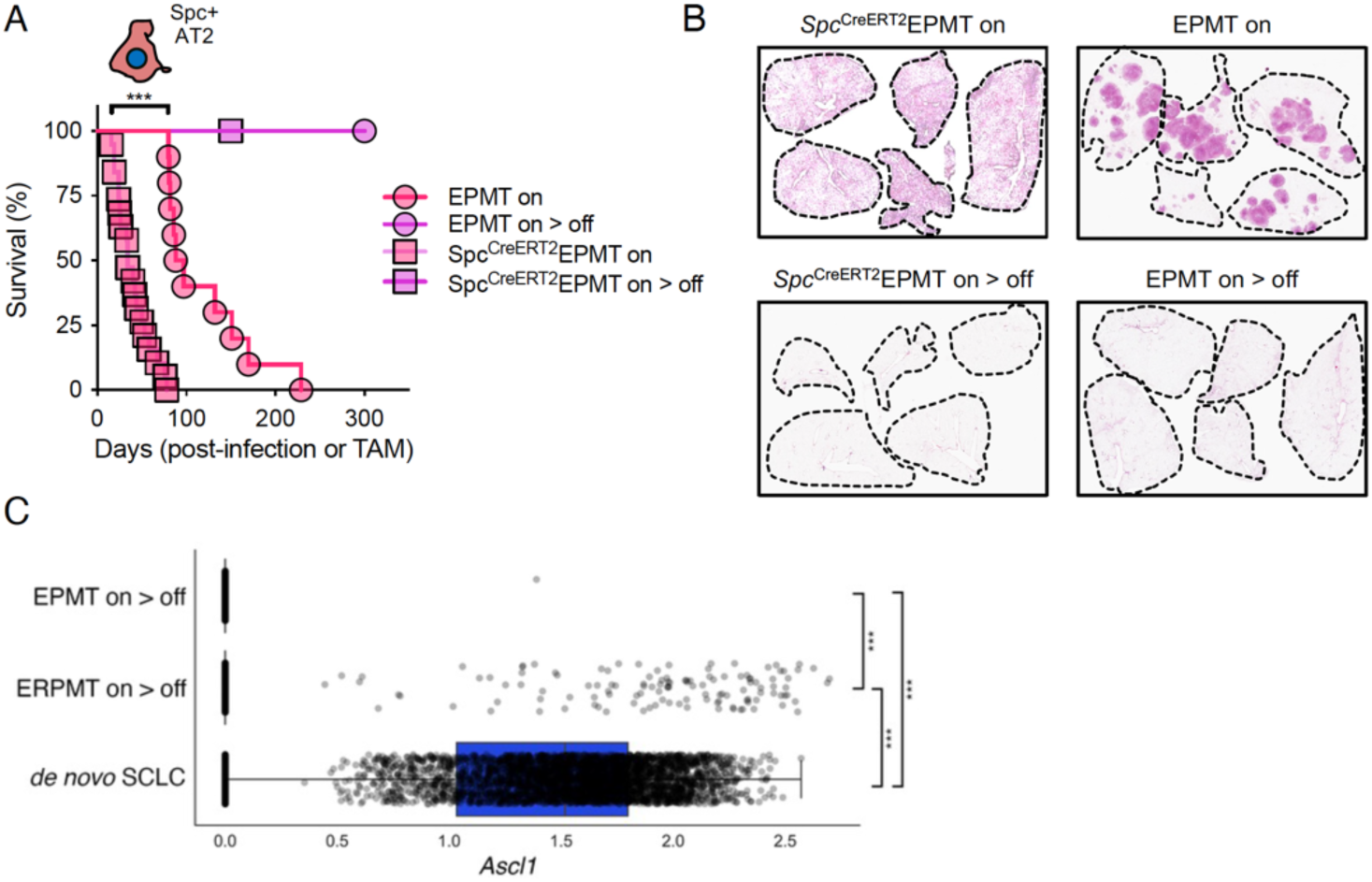
R*b*1 loss is required for the emergence of a neuroendocrine residual from an AT2- derived LUAD. A) Tumorigenesis initiated using an adenoviral (n = 10; Ad5.Spc-Cre) or AT2 lineage trace allele (n = 19; *Spc*^CreERT2^) in the EPMT model (*Rb1*^+/+^) produces LUAD that does not relapse following DOX removal (n = 5 per group). As before, DOX diet was removed when individual mice exhibited signs of labored breathing and/or significant weight loss with a hunched appearance. For each model, cohorts of mice were followed until ∼3X the median latency elapsed, at which point lungs were collected and the study was ended. B) Representative sagittal lung H&E sections from each group in *A* on DOX at point of moribund disease or one month following the removal of DOX from an otherwise moribund animal. C) *Ascl1* expression in single cells isolated from 1mo LUAD residuals in the EPMT and ERPMT models where Cre was expressed in AT2 cells, compared to *de novo* SCLC in the ERPMT model where Cre was expressed in the PNEC lineage, off DOX. Pairwise, multi-hypothesis corrected non-parametric Mann-Whitney U test; p-adjusted < 1e-26 for all (***). 1 of 4,232 cells in the EPMT on > off pool was *Ascl1*-positive and was not a complication of library size, complexity, or poor-quality RNA. We cannot rule out that *Rb1* was not inactivated in this cell.

**Supplemental Figure 15.**
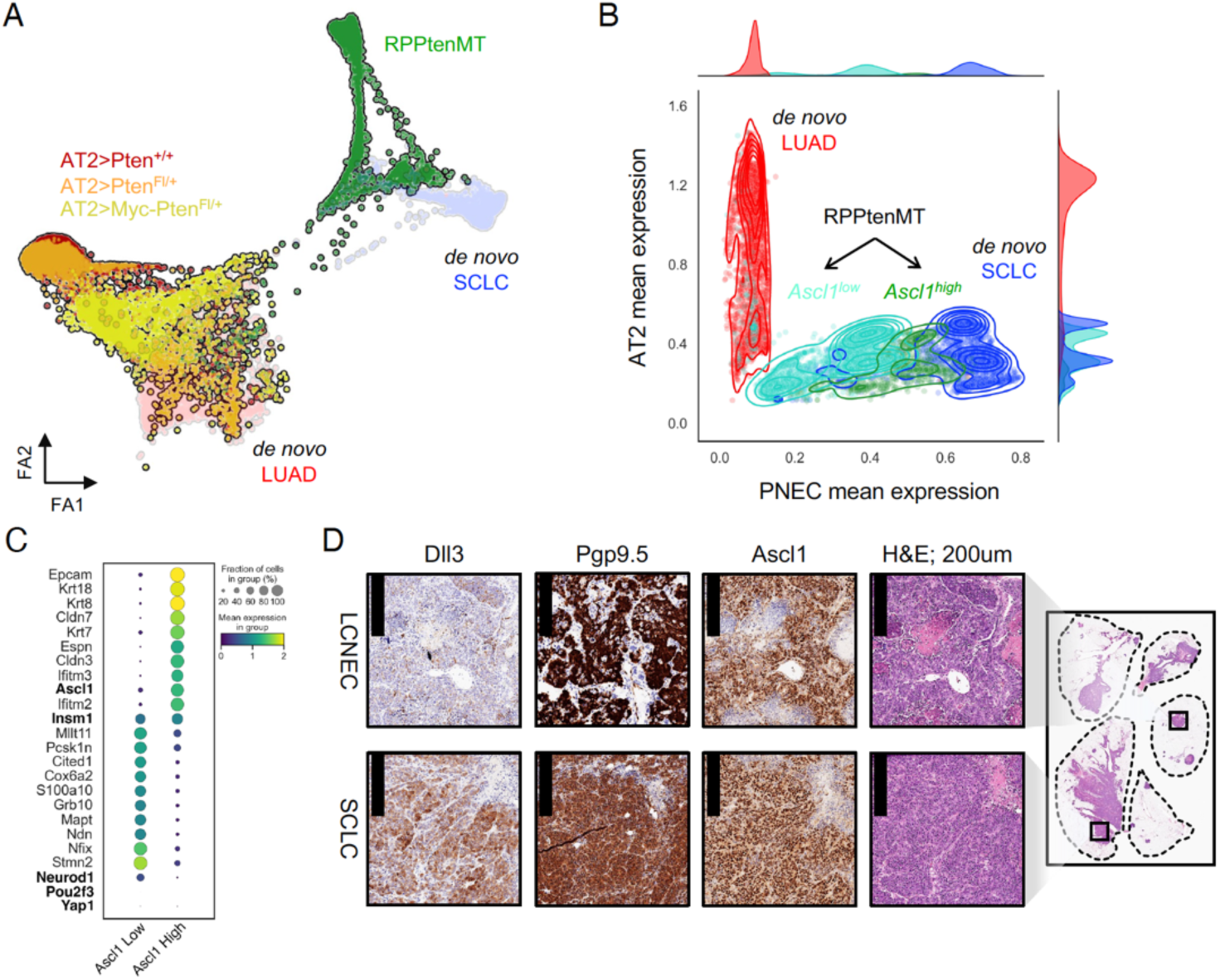
RPPtenMT tumors are composed of *Ascl1^high^* and *Ascl1^low^* cells. A) Force-directed layout of tdTom+ tumor cells sorted from the RPPtenMT model (*green,* n = 5,596 cells) with ERPMT-derived LUAD (*opaque*; *red*, n = 5,394 cells) or SCLC (*opaque*; *blue*, n = 4,371 cells) models with normal (*dark red*, n = 4,429 cells), Pten-deleted (*orange,* n = 4,312 cells), or Pten-deleted and Myc-expressing (*yellow,* n = 3,930 cells) AT2 cells. B) Average imputed expression of AT2 and PNEC lineage markers in tdTom+ cells isolated from ERPMT-derived LUAD (*red*), SCLC (*blue*), and RPPtenMT-derived transformed SCLC (*green*); *Ascl1*-high (*dark green*) and *Ascl1*-low (*teal*) cell populations are highlighted in the RPPtenMT model. C) Dot plot showing frequency of expressing cells (node size) and log-transformed expression (node color) of the top 10 differentially expressed genes (ranked by logFC, Bonferroni p-adjusted < 0.05) between *Ascl1*-high and *Ascl1*-low subpopulations isolated from RPPtenMT tumors (**table S2**). Additionally, SCLC subtype-specific transcription factors are shown bolded, along with *Insm1*, a marker that is occasionally used in the diagnosis of neuroendocrine tumors and expressed here independent of *Ascl1* status. D) Histologic comparison within RPPtenMT tumor-bearing lungs initiated from AT2 cells showing select regions (*boxed*) of large cell neuroendocrine (LCNEC) and pure SCLC histology with indicated neuroendocrine marker staining by IHC identified; 200um scalebar.

